# The impact of PCR duplication on RNAseq data generated using NovaSeq 6000, NovaSeq X, AVITI and G4 sequencers

**DOI:** 10.1101/2023.12.12.571280

**Authors:** Natalia Zajac, Ioannis S Vlachos, Sija Sajibu, Lennart Opitz, Shuoshuo Wang, Sridar V Chittur, Christopher E. Mason, Kevin L Knudtson, John M Ashton, Hubert Rehrauer, Catharine Aquino

**Affiliations:** Functional Genomics Center Zurich, ETH Zurich and University of Zurich, Zurich, Switzerland; Spatial Technologies Unit, HMS Initiative for RNA Medicine, Department of Pathology, Beth Israel Deaconess Medical Center, Boston, MA USA; Cancer Center & Cancer Research Institute, Beth Israel Deaconess Medical Center, Boston, MA, USA; Broad Institute of MIT & Harvard, Cambridge, MA, USA; Center for Functional Genomics, University at Albany, State University of New York, Albany, NY, USA; Iowa Institute of Human Genetics, Genomics Division, University of Iowa, Iowa City, IA, USA; Genomics Research Group, Association of Biomolecular Resource Facilities, USA; Wilmot Cancer Institute, Genomics Research Center, University of Rochester, Rochester, NY, USA; Department of Physiology, Biophysics and Systems Biology, Weill Cornell Medicine, New York, NY, USA; The WorldQuant Initiative for Quantitative Prediction, New York, NY, USA

## Abstract

RNA sequencing (RNA-seq) is a powerful technology for gene expression and functional genomics profiling. Expression profiles generated using this approach can be impacted by the methods utilised for cDNA library generation. Selection of the optimal parameters for each step during the protocol are crucial for acquisition of high-quality data. Polymerase chain reaction (PCR) amplification of transcripts is a common step in many RNA-seq protocols and, if not optimised, high PCR duplicate proportions can be generated, resulting in the inflation of transcript counts and introduction of bias. In this study, we investigate the impact of input amount and PCR cycle number on the PCR duplication rate and on the RNA-seq data quality using a broad range of inputs (1 ng -1,000 ng) for RNA-seq library preparation with unique molecular identifiers (UMIs) and sequencing the data on four different short-read sequencing platforms: Illumina NovaSeq 6000, Illumina NovaSeq X, Element Biosciences AVITI, and Singular Genomics G4. Across all platforms, samples of input amounts greater than 125 ng had a negligible PCR duplication rate and the number of PCR cycles did not have a significant effect on data quality. However, for input amounts lower than 125ng we observed a strong negative correlation between input amount and the proportion of PCR duplicates; between 34% and 96% of reads were discarded via deduplication. Fortunately, UMIs were effective for removing *in silico* PCR duplicates without removing valuable biological information. Removal of PCR duplicates resulted in more comparable gene expression obtained from the different PCR cycles. Data generated with each of the four sequencing platforms presented similar associations between starting material amount and the number of PCR cycles on PCR duplicates, a similar number of genes detected, and comparable gene expression profiles. However, the sequencers using conversion kits for Illumina libraries (AVITI, G4) exhibited lower adapter dimer abundance across all input amounts, but also a higher PCR duplication rate in very low input amounts (<15ng). Overall, this study showed that the choice of input amount and number of PCR cycles are important parameters for obtaining high-quality RNA-seq data across all sequencing platforms. UMI deduplication is an effective way to remove PCR duplicates, improving the data quality and removing any variation caused by the conversion kits.

## Introduction

RNA-seq is a technology applied for quantification of RNA abundance and allows the study of gene regulation and function (Hrdlickova et al., 2017). This approach has been widely applied from quantification of expression of all genes across tissues or cells to the targeted quantification of specific species of transcripts (Z. Wang et al., 2009). Prior to sequencing, extracted RNA is converted to cDNA and during library construction, it is amplified via polymerase chain reaction (PCR) to enrich properly structured fragments bearing ligated adapters and generate adequate input material for sequencing. PCR amplification is known to introduce bias due to unequal probabilities of amplification of certain molecules, which in turn can impact the accuracy, sensitivity and precision of transcript quantification (Parekh et al., 2016). The amount of input material and the number of PCR cycles directly impact the proportion of spurious duplicate reads but the optimal set of parameters depends on the library complexity and sequencing depth (Fu et al., 2018).

In DNA sequencing, duplicates can be identified and marked with tools such as Picard (MarkDuplicates), where they are identified based on the alignment coordinates of the 5′ end of each read (Faust & Hall, 2014), and then subsequently flagged or removed in downstream analyses. However, in RNA-seq, the distinction of amplification-derived duplicates cannot be performed *in silico* purely by mapping coordinates (X. Li et al., 2015; Picelli et al., 2014) and could often result in removal of a large proportion of biologically relevant information (Sayols et al., 2016). To this end, Unique Molecular Identifiers (UMIs), which are short (often 5-11 nucleotides) random stretches of oligonucleotides, can be added to the RNA fragments prior to amplification (Fu et al., 2018; Hrdlickova et al., 2017; Parekh et al., 2016), to enable the detection of individual molecules. Following sequencing, a computational model accounting for UMI errors can be applied to identify reads with identical alignment coordinates and identical UMI sequences which are then assumed to be duplicates (Fu et al., 2018; Islam et al., 2014; Smith et al., 2017).

The most widely adopted short-read sequencing technology used for RNA sequencing is Illumina sequencing by synthesis (SBS), also called the Illumina dye sequencing (Ambardar et al., 2016), although a range of other long-read sequencing platforms have also been used for these types of experiments since 2010 (Cho et al., 2014; S. Li et al., 2014). Other short-read technologies and instruments have been recently introduced to the market, proposing alternative sequencing approaches which could provide specific improvements in cost, flexibility, sequencing time, and/or throughput (Eisenstein, 2023; LeMieux, 2022), including the G4 and AVITI benchtop sequencers, from Singular Genomics and Element Biosciences, respectively. The former combines short sequencing times with 4 flow cells in parallel, allowing pooling of experiments and improving efficiency, while the latter uses the alternative sequencing by binding (SBB) chemistry that involves the binding of a multivalent fluorescent polymerase substrate by avidity which is suggested to improve read accuracy and reduce costs (Arslan et al., 2023; Biswas et al., 2022).

There are currently multiple different vendors providing specific library preparations for all aforementioned technologies. However, the need of sequencing an RNA-seq library generated using Illumina-specific reagents on alternative sequencers is still a common scenario. These libraries contain Illumina specific adapters, which need to be converted prior to sequencing on a different platform. The conversion protocols include additional PCR steps, which could potentially introduce additional biases such as an increase in the rate of PCR duplicates (Eisenstein, 2023).

In this study, we performed bulk RNA sequencing of a human liver RNA sample, creating serial dilutions of input amounts ranging from 1 ng to 1000 ng (Fig.1). Illumina libraries were prepared using a strand-specific protocol, able to support starting material of 10 ng to 1,000 ng of RNA (New England Biosciences). Unique Dual Index UMI adapters were incorporated to mark unique fragments, and each input amount was amplified with three different numbers of PCR cycles (Low, Intermediate, High, with a 2-cycle increment between the different levels). The native Illumina libraries were directly sequenced on two Illumina sequencers, a NovaSeq 6000 and a NovaSeq X. Following conversion protocols from the vendors, the same pooled Illumina libraries were sequenced on an Element Biosciences AVITI system, and a Singular Genomics G4 sequencer. Our study aimed to address the rate of PCR duplicates in bulk RNA sequencing data focusing on how the rate of PCR duplication is affected by 1) the amount of input material and number of PCR cycles and 2) the conversion process employed to render an Illumina library compatible with the new short-read sequencing technologies. We also evaluated the quality of the data addressing potential biases introduced by the amount of input material and the number of PCR cycles on the number and nucleotide composition of the detected genes and gene expression. Importantly, we offer an outlook of the new sequencing instruments (Illumina NovaSeq X, AVITI, G4) focusing on instrument-specific duplication phenotypes, as compared to the widely deployed NovaSeq 6000 platform.

**Figure 1.**
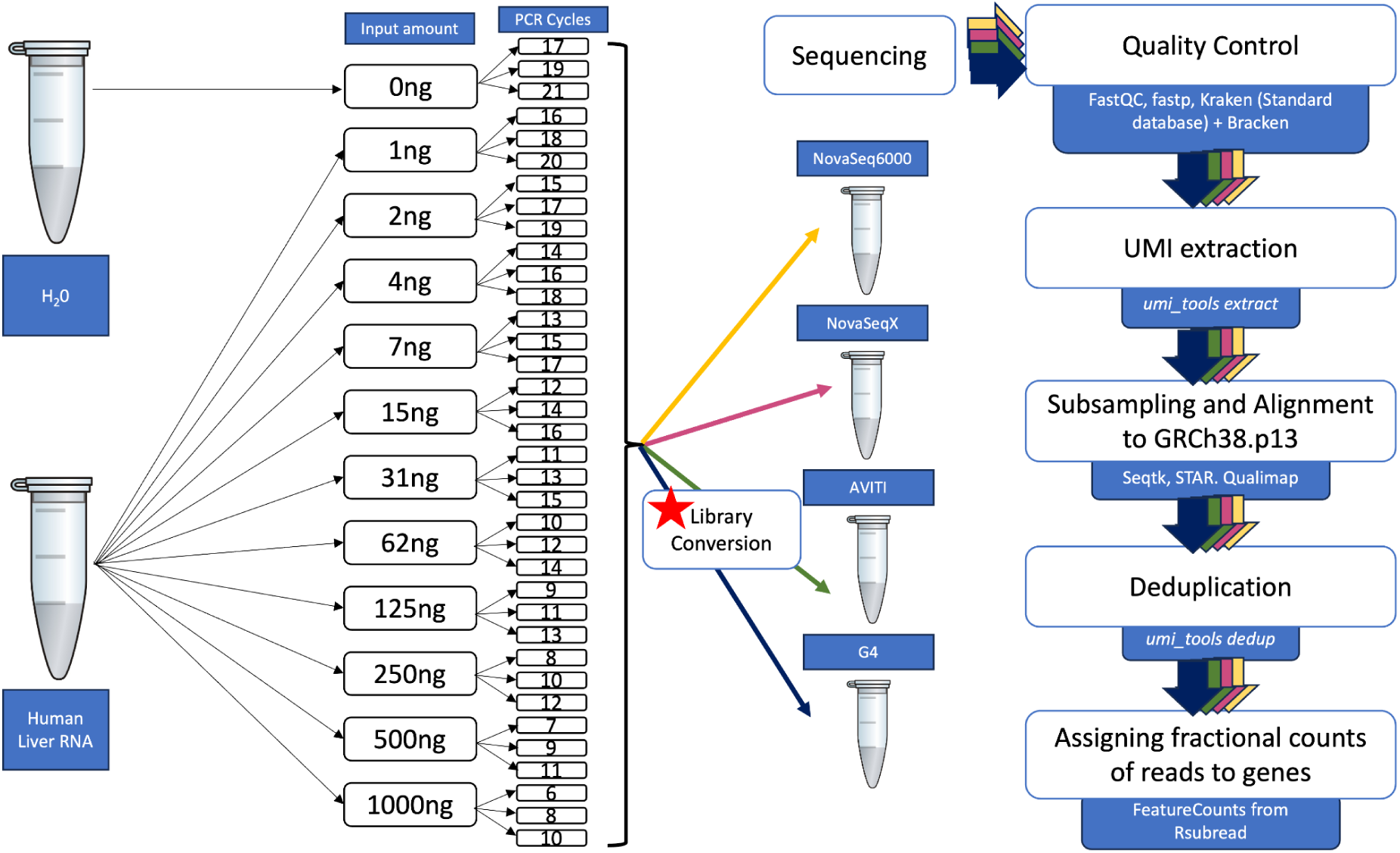
Summarised experimental workflow.

## Methods

### Library construction

A serial dilution from 1 ng to 1,000 ng Human Liver Total RNA (ThermoFisher) was prepared to generate the various input samples. The NEBNext Ultra II Directional RNA Library Prep Kit for Illumina with Unique Dual Index UMI Adapters RNA Set1 (NEB, Franklin Lake, NJ, USA) were used in the succeeding steps according to manufacturer’s instructions. Briefly, total RNA samples (1 ng -1,000 ng) were polyA enriched and then fragmented prior to reverse-transcription into double-stranded cDNA. The cDNA samples were end-repaired before ligation of adapters containing UMI. Fragments containing adapters on both ends were selectively enriched with PCR containing unique dual indices (UDI) for multiplexing. Per dilution, 3 different PCR cycles were used. The quality and quantity of the enriched libraries were validated using a TapeStation (Agilent, Santa Clara, California, USA). The product is a smear with an average fragment size of approximately 260 bp. The libraries were normalised to 10nM in Tris-Cl 10 mM, pH8.5 with 0.1% Tween 20. As the different dilutions and PCR cycles used resulted in very varied library concentrations, the pooling was simplified by using 5 ul of the libraries produced.

### Next Generation Sequencing

#### Illumina NovaSeq 6000 and Illumina NovaSeq X

The pool of Illumina libraries was quantified using a TapeStation (Agilent, Santa Clara, California, USA) and normalised to a loading concentration specific for the instrument type. For the NovaSeq 6000 (Illumina, Inc, California, USA), 18ul of the pooled libraries with a concentration of 0.8 nM was loaded on a lane of a NovaSeq 6000 SP Reagent Kit v1.5 (100 cycles) flowcell for a final loading concentration of 180 pM. For the NovaSeq X (Illumina, Inc, California, USA), 34uL of the pooled libraries with a concentration of 0.55 nM were loaded into a lane of a NovaSeq X Series 10B Reagent Kit (300 Cycle) flowcell for a final loading concentration of 110 pM. The pools were sequenced single-end 100 bp on the NovaSeq 6000 and paired-end 150 on the NovaSeq X.

#### Element Biosciences AVITI

The pool of Illumina libraries was prepared for sequencing on the AVITI sequencer (Element Biosciences, San Diego, CA) using the Element Adept Library Compatibility Kit v1.1. This process involves the denaturation, library circularization via ligation to a splint adapter, and exonuclease digestion of non-circularized molecules. Thirty microliters of the Illumina sequencing library pool at a concentration of 16.7 nM were circularized. The resulting circularized library was quantified via qPCR using the standards provided in the compatibility kit and qPCR (SYBR Green PCR Master Mix, Applied Biosystems, Waltham, MA). Twenty-five microliters of the circularized library at a concentration of 3.5 pM were loaded onto the AVITI system and sequenced with a AVITI 2x150 Sequencing Kit.

#### Singular Genomics G4

To enable anchoring of clusters on Singular flow cells, custom conversion primers targeting P5 and P7 with Singular specific (S1 and S2) 5’ overhangs were used to retain the original indexes. For sequencing, 2mL of custom index primers (1uM) were loaded into the custom primer wells of 300 Cycle reagent cartridges (Lot 2304251). The library pool was quantified with a dsDNA HS Assay Kit (Q33230) on a Qubit 4 fluorometer (ThermoFisher) and diluted down to 1ng/uL with Ambion RNAse-free water, then amplified with Roche KAPA HiFi HotStart ReadyMix in a BioRad C1000 thermal cycler for a total cycle number of 7. Annealing was set for 30 seconds at 57°C. Subsequently, PCR product was cleaned up with SPRISelect beads (Beckman-Coulter) and verified on Fragment Analyzer 5200 using Agilent DNF-473 NGS Fragment Kit (1-6000 bp). To determine the optimal loading concentration, 200 pM libraries and the 50 pM PhiX Control were diluted down to perform a titration run with a series of 12 pM, 15 pM, 17 pM and 20 pM. Final sequencing was performed using 15 pM loading concentration on two separate F2 flow cells (Lot 4052120, Serial number OM0075H and #OM0075H) using a read length setup of 8 (i5, index 1): 100 (Read 1) as well as 19 (i7, index 2 and UMI): 100 (Read 2). Noteworthy, on the Singular G4 platform, i5 corresponds to index 1 and i7 corresponds to index 2, instead of reverse complement i7 and i5, respectively, as on the Illumina or AVITI platform.

### Demultiplexing

The demultiplexing for the G4 and Aviti data was performed with the sgdemux tool (https://github.com/Singular-Genomics/singular-demux). The Illumina data were demultiplexed using bcl2fastq v2.20

(https://support.illumina.com/downloads/bcl2fastq-conversion-software-v2-20.html). The UMIs were located within the i7 adapter sequence in either position 1 to 11, as for NovaSeq 6000 and NovaSeq X data, or 2 to 12, as for AVITI and G4 data.

### Bioinformatics

The quality of the data was assessed using FastQC v0.11.9. For each sequencing technology, the data were subsampled to 2,000,000 reads per sample (or maximum number of reads if the number of reads was lower than 2,000,000). Even though the sequencing was performed in a paired-end mode using G4, AVITI and NovaSeq X, only the forward reads were used in the analysis for even comparison with the NovaSeq 6000 sequencing, which was performed in single-end mode. The subsampled reads were processed using fastp v0.23.4, which involved trimming of Illumina adapters and filtering of reads below 18 bp in length (Chen et al., 2018). NovaSeq X reads were trimmed to the length of 101 bp to match the length of the reads from the other sequencers. The number of reads filtered due to length was used for estimation of the proportion of primer dimers in each sample. Subsequently, the level of contamination in the filtered reads was estimated with Kraken2 v2.0.9 using the Standard database (05.06.2023) followed by Bracken v2.8 for abundance estimation of human and non-human reads (Lu et al., 2017, 2022; Wood et al., 2019; Wood & Salzberg, 2014). Abundance estimation was run at a genus level. The final abundance was re-estimated after including reads unclassified by Kraken.

The data were then processed using UMI-tools and STAR (Dobin & Gingeras, 2016; Smith et al., 2017). First, the UMIs were extracted from the reads and inserted into the read names using the *umi_tools extract* option. The reads were then mapped with STAR in a 1-pass mode to GRCh38.p13 reference (Gencode 42 release), allowing a maximum of 10 mismatches between the read and the reference and a maximum of 50 multiple alignments per read, outputting alignments only if the number of matched bases was higher than 30bp. Deduplication was performed with *umi_tools dedup*, using the directional method for identifying clusters of connected UMIs and an edit distance threshold of 1. Number of aligned reads was obtained by computing the number of primary alignments using samtools flagstat (samtools v1.17). The mapping quality (mismatch rate and GC content) of the raw and deduplicated data was assessed with Qualimap v2.2.1 (García-Alcalde et al., 2012; Okonechnikov et al., 2016). FeatureCounts from the Rsubread v2.14.1 Bioconductor package (Liao et al., 2014) was employed for assigning mapped sequencing reads to genomic features taking into account the multi-overlapping and the multi-mapping reads, counting each alignment fractionally. R packages such as stats v4.3.0, umap v0.2.10.0 and rstatix v0.7.2 were respectively used for performing Pearson’s correlation, the UMAP analysis and carrying out Friedman’s test followed by Wilcoxon signed-rank test. The UMAP was done on normalised and log2 transformed counts with an added background expression of 1. The counts were normalised using the scaling normalisation method from EdgeR.

All scripts are included in Supplementary Information.

## Results

### Featured datasets

We generated complex libraries from a human tissue sample (human liver RNA) at different dilutions (1ng to 1000 ng) and 3 different levels of amplification (categorized as Low, Intermediate, High, with a 2 cycle difference between consecutive levels). The number of cycles was adjusted to the input amount, as recommended in the manual (New England Biolabs, 2019) (Fig. 1). Following a multi-center setup, the samples were sequenced on four sequencers in three different laboratories and sequencing facilities, the Functional Genomics Center Zurich of the ETH Zurich and University of Zurich, DNA Technologies and Expression Analysis Core, UC Davis Genome Center at UC Davis, and the Spatial Technologies Unit of the Harvard Medical School Initiative for RNA Medicine at Beth Israel Deaconess Medical Center. The data from all four sequencers were subjected to the same bioinformatics workflow following customised sample demultiplexing, performed according to sequencer manufacturer guidelines (Fig. 1).

### Analysis of raw reads reveals that evaluation of sample quality is required prior to sequencing

The study setup enabled us to extract in-depth information about data quality, quantity, and complexity across the different sequencing platforms, including Phred quality scores across the read length, sequence biases, duplication rate, and adapter primer dimer contamination.

The produced reads were 100 bp long for G4 and Aviti, 101 bp long for NovaSeq 6000, and 151 bp long for the NovaSeq X, subsequently also trimmed to 101 bp. In terms of sequencing accuracy scores, the average Phred quality score of reads across all sequencers was high and ranged from 36 to 43, highest for reads sequenced with the AVITI (Supplementary Fig. 1). Adapter dimer contamination was one of the few metrics which has highly differentiated between the different platforms. For NovaSeq X and NovaSeq 6000, the adapter dimer contamination, or the number of reads discarded due to not meeting length criteria (<18bp), was higher than for the AVITI and G4 sequencers. (Fig. 2). The percentage of primer dimers for the Illumina sequencers ranged from 5.6% to 70.1% for samples of input amounts between 1 ng and 15 ng and from 1.3% to 16.6% for input amounts above 31 ng, with a 10% to 25% increase from NovaSeq 6000 to the NovaSeq X. Two samples of input amounts 250 ng and 500 ng, amplified using the highest value of PCR cycles and sequenced with the Illumina sequencers, exhibited a fraction of primer dimers comparable to that of the low input amounts (between 41% - 62%), thus creating outliers. Both the AVITI and G4 exhibited low primer-dimer amounts, which can be attributed to the additional library conversion steps and library cleanup. The percentage of adapter primer dimer contamination for the G4 and AVITI ranged from 0.0088% to 3.3% across all input amounts (1 ng -1,000 ng) and went up to 11.1% in the negative control (Fig. 2).

**Figure 2.**
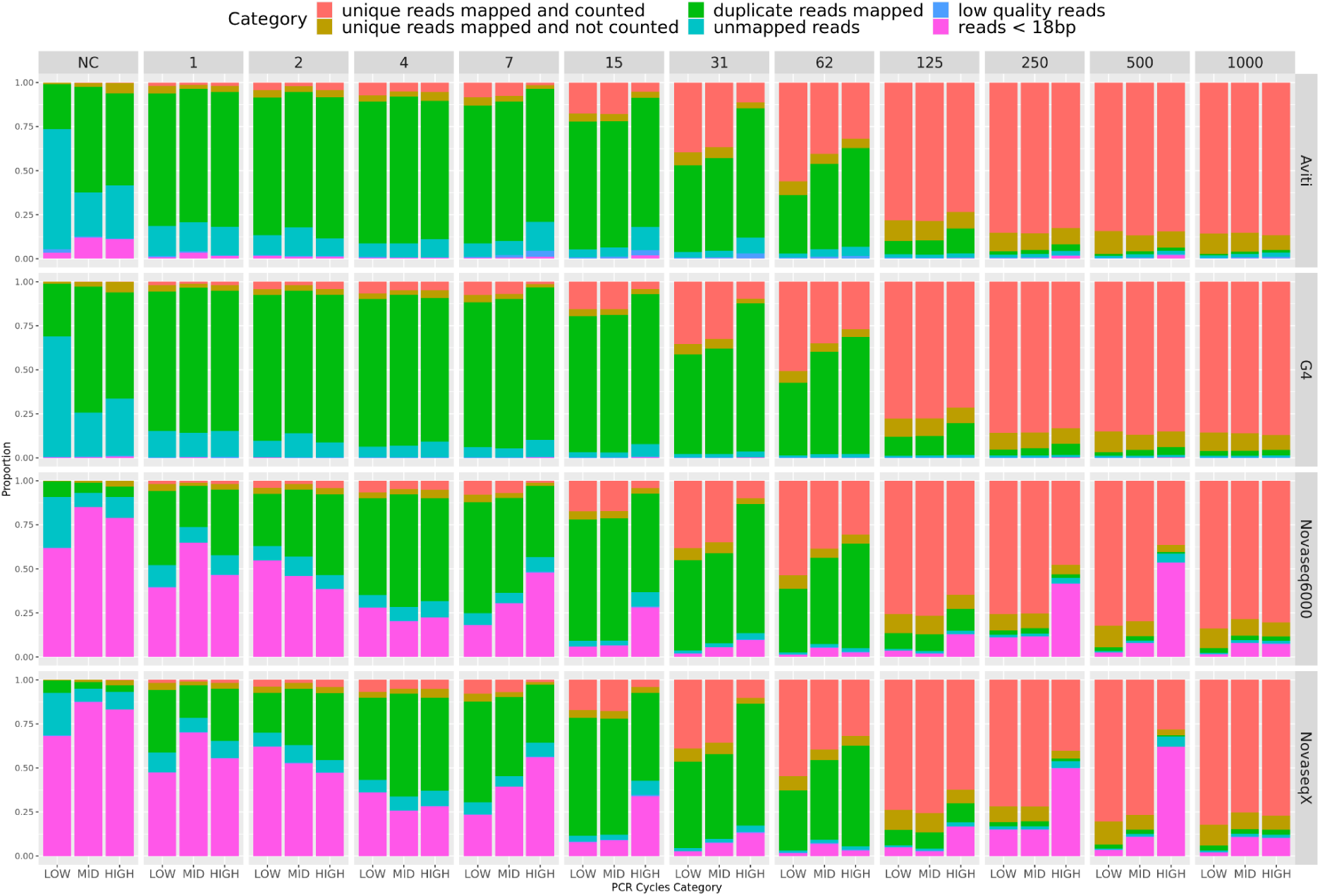
The classification of total subsampled reads per each sample. The colors indicate the categories: low quality and reads shorter than 18bp were filtered with fastp, unmapped reads were rejected by the STAR mapper, duplicate reads were removed via deduplication and only mapped and counted unique reads represent gene expression. Counts used for this figure are attached as Supplementary Table 1. Data sequenced with NovaSeq X and NovaSeq 6000 had a considerably higher proportion of short and low quality reads/ contamination with adapter primer dimers than the data sequenced on AVITI and G4. AVITI and G4 exhibited a higher proportion of PCR duplicates for low input amounts. Adapter primer dimers were cleaned up from the AVITI and G4 data during library cleanup performed during conversion.

Another important feature in data quality is the sequencing error rate. The most common type of error in Illumina sequencing data are mismatches, with a median of 0.1 per base for NovaSeq 6000 (Stoler & Nekrutenko, 2021). Thus we assessed the error rate across all samples using the rate of mismatches in raw reads mapped to the human genome. We observed a low rate of mismatches for all samples with input amount greater than 1 ng, varying between 0.0003 and 0.001. The rate of mismatches decreased with increasing input amount and for input amounts between 4 ng and 31 ng there was an elevation of the rate of mismatches for the highest PCR cycle category, suggesting introduction of errors during amplification (Fig 3, Supplementary Table 2). We observed no difference in mismatch rate between NovaSeq X, NovaSeq 6000 and AVITI but the data sequenced with G4 had an approximately 50% increase in mismatch rate compared to the other sequencers (Fig 3).

**Figure 3.**
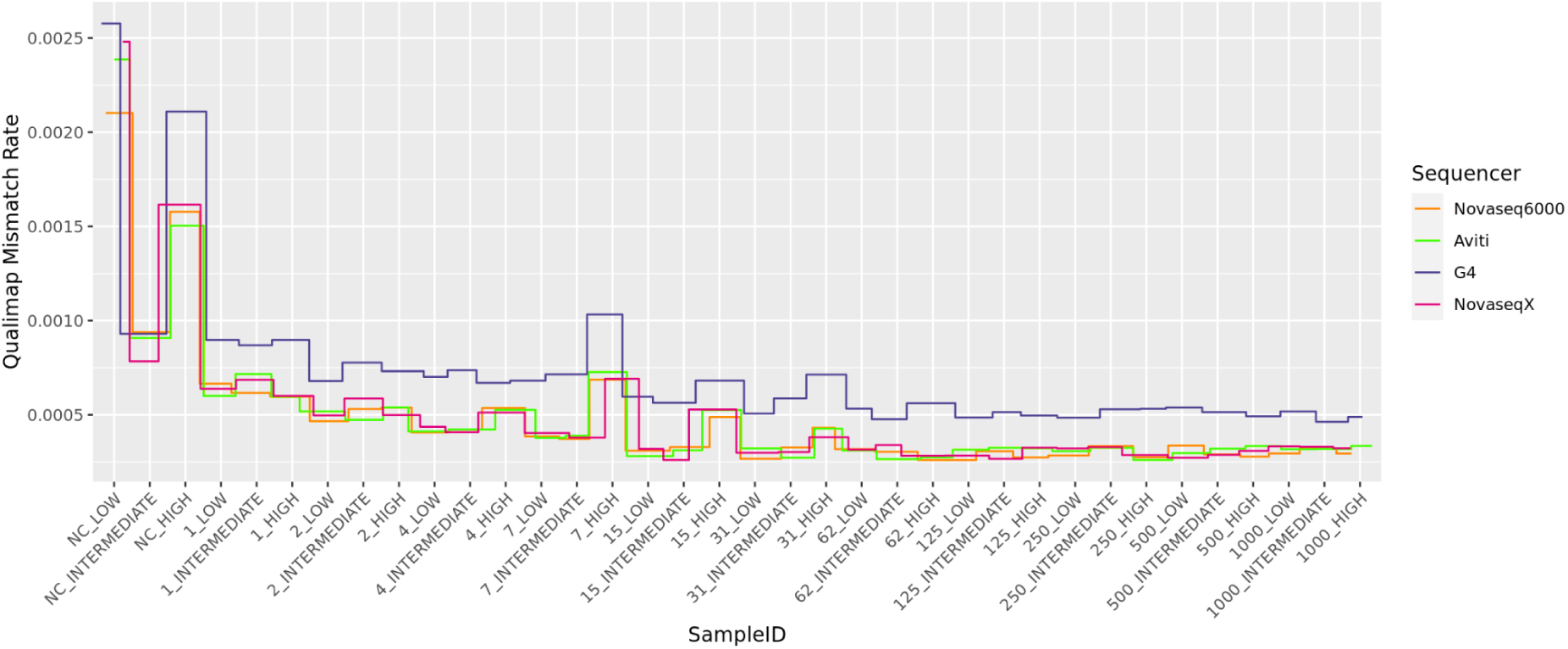
Rate of mismatches in raw data mapped to the human genome, measured with Qualimap (v2.2.1). The generally low mismatch rate decreased with increasing input amount. For low input amounts, there was an elevated rate of mismatches in the highest PCR cycle category. G4 had a higher rate of mismatches compared to the other sequencers.

RNA samples often contain a small fraction of microbial contamination but elevated microbial content can be another reason for considerable data loss during preprocessing and can indicate potential issues in handling of samples, such as contamination of reagents (de Goffau et al., 2018). We assessed contamination in all samples after filtering of the data for short and low quality reads (after removal of the reads marked in pink on Fig 2). As expected, we found the highest fraction of microbial contamination in negative control samples (0 ng of input amount) and with similar proportions across the different sequencers (Fig 4, Supplementary Table 3). For the NovaSeq X and NovaSeq 6000, 28% and 31% of the reads, respectively, were identified *in silico* as of bacterial origin, 35%-41% were unclassified, and 31-34% of the reads were human RNA. The majority of the bacterial reads mapped to *Cutibacterium, Streptococcus, Staphylococcus and Pseudomonas*. For the AVITI and the G4, the 0 ng sample had a similar contamination profile with 60%-65% of the reads mapping to human and 22%-24% being of non-human origin, in majority also mapping to *Cutibacterium, Streptococcus, Staphylococcus, and Pseudomonas.* Only 12-16% of the reads were unclassified. For samples between 1 ng and 15 ng the microbial contamination ranged from 8% to 1.5% respectively across all sequencers. Samples of input amount above 31 ng consisted only of human and unclassified reads with insignificant traces of microbial content. For the NovaSeq 6000 and NovaSeq X, the proportion of unclassified reads appeared to increase again to 2.7% for samples at 250 ng and 500 ng, where the highest proportion of primer dimers were identified prior to filtering.

**Figure 4.**
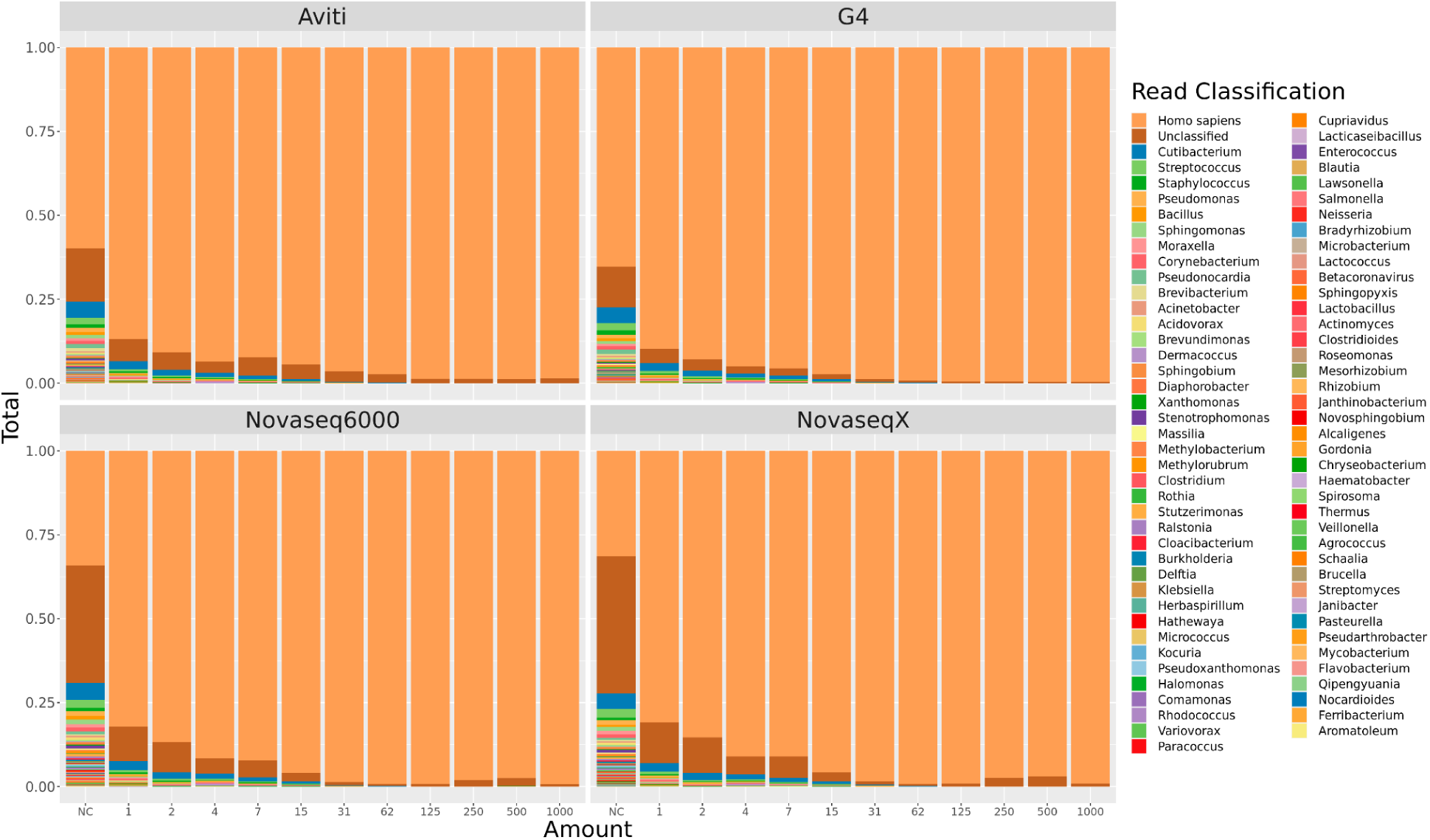
Taxonomic classification, assessed with Kraken and Braken, of the subsampled reads, filtered of low quality and short reads. The data is displayed as per sequencer per input amount and the result is an average across all three PCR cycles. The data is sorted by read abundance. The first two categories represent reads mapping to the human genome and reads unclassified by Kraken standard database (05.06.2023). Highest proportion of non-human RNA contamination was found in the negative control samples and samples of input amount below 7 ng.

### Number of artifactual reads depends on a combined effect of input amount and the number of PCR cycles

NEBNext Ultra II Directional RNA Library Prep Kit for Illumina requires between 10 ng and 1,000 ng of input material (New England Biolabs, 2019). To test whether the input amount or the value of PCR cycles chosen for amplification had a significant effect on the rate of PCR duplicates, and thus the recovery of usable reads, we evaluated the percentage of aligned reads in the raw data and after deduplication (Fig. 2). The percentage of mapped reads reported here was calculated as the number of aligned raw and unique reads relative to the total number of subsampled reads for each sample (see Fig. 5 for the total number of subsampled reads). Additionally, the percentage of PCR duplicates was computed as the percentage of aligned reads that was removed by the deduplication algorithm of *UMI-tools*.

**Figure 5.**
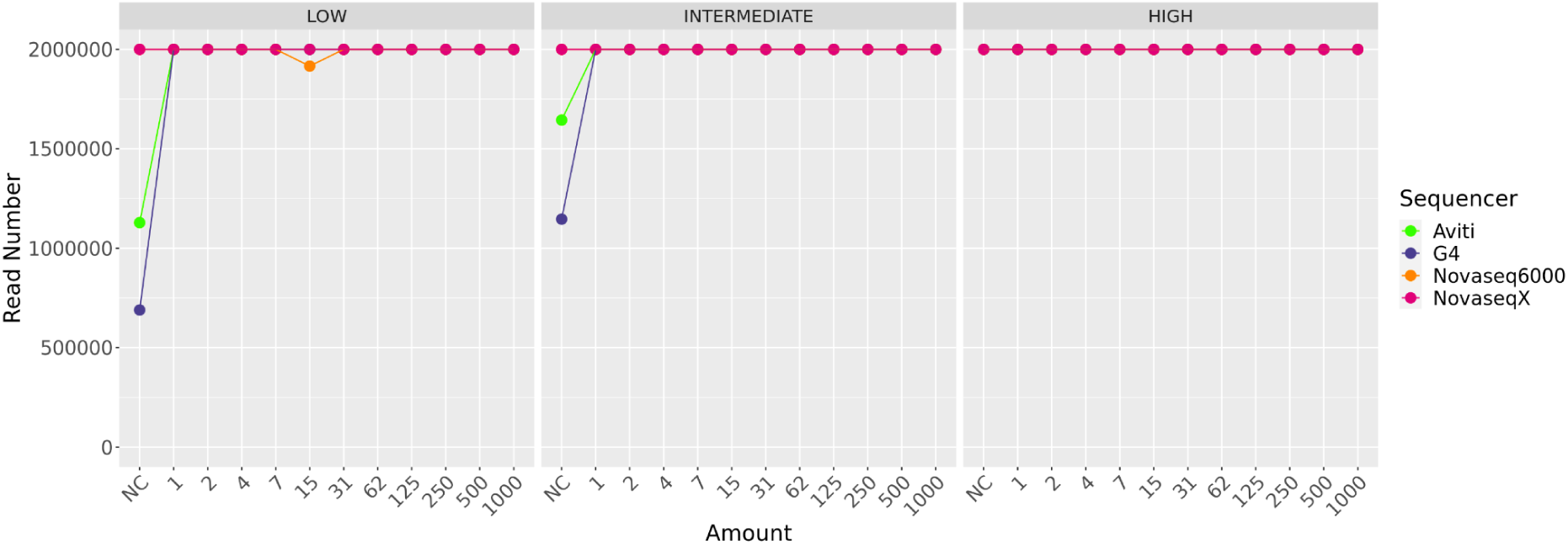
The total number of subsampled reads per each sample.

The percentage of aligned reads before and after deduplication generally increased with input amount across all 3 PCR cycle categories (Fig 6). For low input amounts, i.e. between 1 ng and 15 ng, there was a clear discrepancy in the aligned raw reads between the four sequencers. For NovaSeq X and NovaSeq 6000, the percentage of mapping reads varied between 21.5% and 91%. Whereas for the AVITI and G4, the values ranged between 79% and 97%. For both AVITI and G4, fewer reads were discarded due to low quality or short length but more reads were identified as a result of PCR duplication. The discrepancy between the sequencers disappeared entirely after removal of duplicate reads; the percentage of reads that mapped to the genome for all sequencers ranged between 3% and 22%.

**Figure 6.**
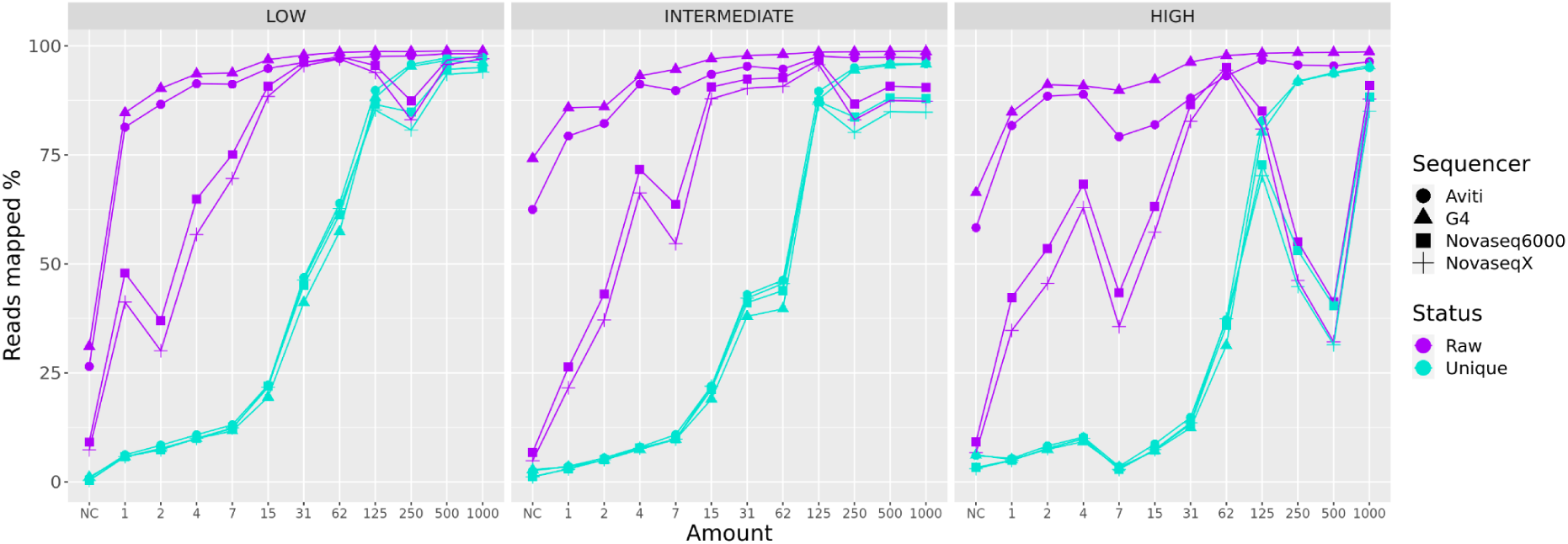
The percentage of reads per sample mapping to the human genome, calculated as the number of reads out of all the subsampled reads. The data is divided by PCR cycle category and the colour indicates whether the data is before (purple) or after (turquoise) deduplication. The difference in the percent of mapped reads between the raw and the deduplicated data was high for input amounts below 125 ng and for samples amplified with the highest PCR cycle category. AVITI and G4 exhibited higher proportion of duplicates than Illumina sequencers for low input amount (below 15 ng).

For the lowest input amount of 1 ng, between 2.8% and 6.2% of the reads were retained after removal of duplicate reads; the highest fraction of productive reads was obtained for the lowest PCR cycle category for all sequencers (5.6%-6.2%). The same was observed for the input amounts between 7 ng and 125 ng; the percentage of duplicates was the highest for the highest PCR cycle category, varying between 82% and 96% for the amount of 7 ng and between 8% and 18% for the 125 ng input. Increasing the number of PCR cycles from Low to Intermediate did not result in a significant increase in the PCR duplicates, except for input amount of 62 ng, with a shift from 34%-42% of duplicates to 50%-60% from the Low to Intermediate PCR cycle category (Fig 2 and Fig 9.Top).

Using input amounts above 250 ng yielded the highest mapping rate and the highest proportion of retained reads, and the results were not influenced by the value of PCR cycles used for amplification (Fig 6). The percentage of discarded reads ranged from 0.8% to 6.6% and the percentage of productive reads retained after deduplication ranged between 85% and 97%. Sample differences stemmed rather from technical issues during library preparation. For 250 ng and 500 ng samples sequenced with NovaSeq 6000 and NovaSeq X, the number of mapped reads amounted to less than 53% in the highest PCR cycle category and between 80% and 88% in the intermediate cycle category. The read dropout can be explained by the high percentage of primer dimers in both samples (between 41% -62%), most noticeably produced when the highest number of PCR cycles was applied.

For the negative control (0 ng of input material), we were able to obtain 2 million reads from all sequencers at the highest cycle category but for intermediate and low PCR cycle categories, we obtained much fewer reads from AVITI and G4 (Fig. 5). After the data were deduplicated, the number of reads remaining for each sequencer in the negative control ranged from 6,301 to 65,838, for NovaSeq X and NovaSeq 6000, from 7,532 to 123,851 for G4 and from 9,503 to 120,954 for AVITI, accounted for by the cross contamination with human RNA. We compared the distribution of all alignments from the negative control to that of samples of 1,000 ng input and observed that most of the reads from the negative control mapped to lncRNAs (16.6%-21.3%), mRNA introns (24.6%-30%), and unannotated regions (43%-52%). Only 30% to 38% of alignments were to coding regions and even a lower percentage, only 3%-13%, belonged to exons (Fig 7). Thus, the unique reads counted into gene expression were those remaining after filtering of non-coding material and that ranged from 142 to 2,738 of reads for NovaSeq X, NovaSeq 6000 and G4 and between 1,530 and 5,658 for AVITI.

**Figure 7.**
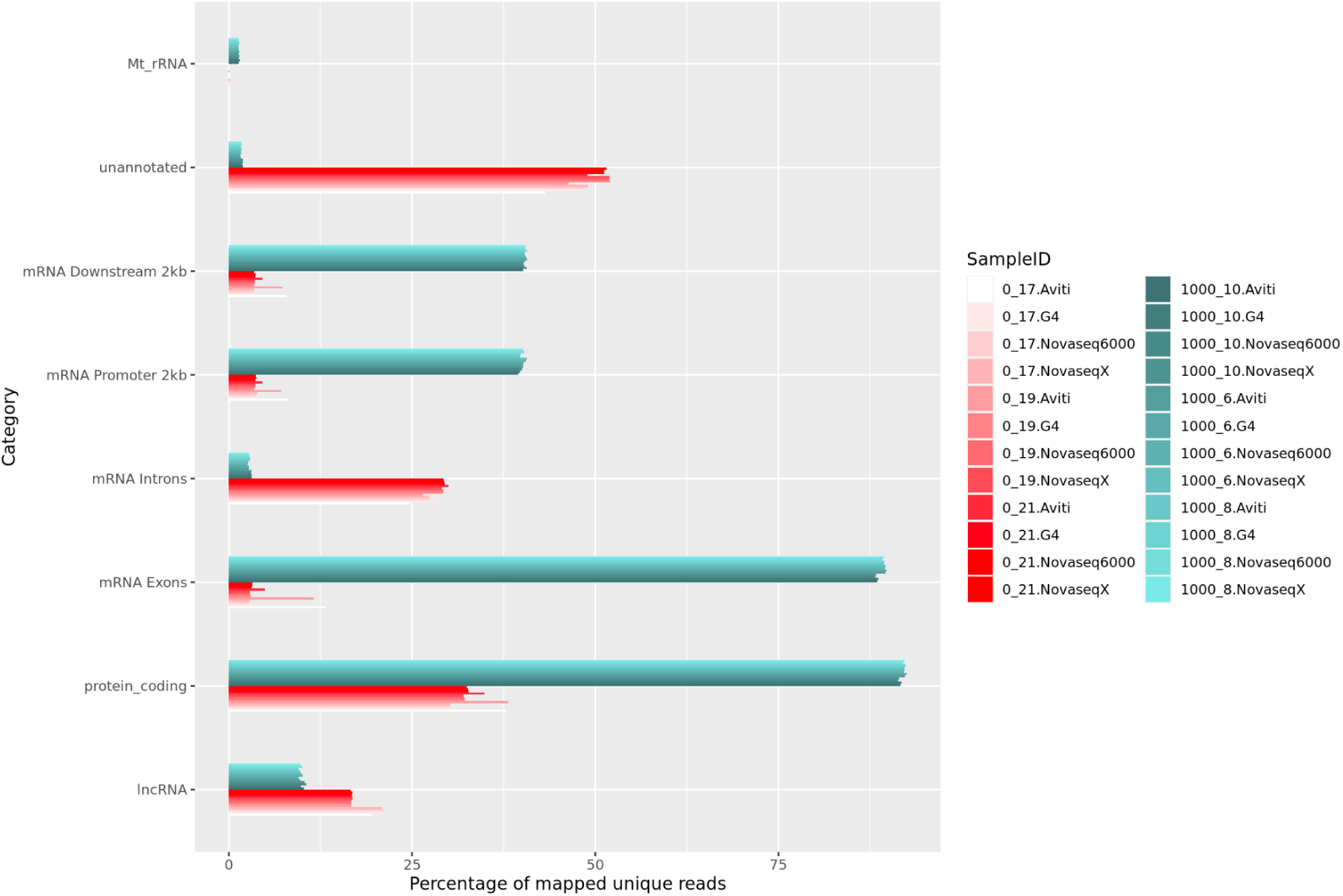
Proportion of alignments to different features, counting all alignments in the deduplicated bam files (Supplementary Table 4). Here we compare the negative control samples (0 ng) to the 1,000 ng samples that had the most uniform coverage across the genome. The human RNA contamination from the negative control samples mapped unevenly across the genome, mostly to unannotated and intronic regions and long non-coding RNA.

### Number of detected genes is positively correlated with the input amount and can be obscured by the rate of PCR duplicates

The main goal of most RNA-seq analyses is to study the gene expression within a sample or to compare the relative gene expression between samples/groups. For low input amounts, even with the best library protocols, there is a higher chance of loss of information with the loss of input material during sample processing (J. Wang et al., 2019). Additionally, low input amounts require higher amplification for obtaining sufficient material for sequencing, leading to a higher tendency for highly expressed genes to produce identical fragments, which in turn can lead to lower probability of sampling the transcripts of lowly expressed genes during sequencing (Fu et al., 2018). PCR duplicates create noise that increases the false positive rate, obscuring the number of detected genes and interfering with the absolute and relative quantification of expression (Parekh et al., 2016; Sheng et al., 2017). To assess the impact of input amount and rate of amplification on gene expression results, for each sample we estimated the number of detected genes as well as gene-wise base expression before and after deduplication.

We found that the number of detected genes positively correlated with the input amount (Fig 8). The number of genes ranged from 5,013 at 1 ng to 14,536 at 1,000 ng, across all sequencers. For input amounts above 125 ng, there was no increase in the number of detected genes (Fig 8). We also found that the number of PCR cycles had an effect. For all sequencers, the lowest number of genes was observed at the highest PCR cycle category for input amounts between 7 ng and 62 ng (Fig 8). At 7 ng input amount, around 50% more genes were detected when the data was amplified with the Intermediate or Low number of PCR cycles as compared to the High number of PCR cycles. For 62 ng that difference decreased to only 5% more genes. For those samples there was a clear correlation between the percentage of PCR duplicates and the number of detected genes (Fig. 9). For input amounts above 125 ng, we observed no effect of the PCR cycles category except for two outliers: 250 ng and 500 ng, where between 13,063 and 13,828 genes were detected for the highest PCR cycles category. Otherwise, the number of genes detected in input amounts above 125 ng ranged between 14,226 and 14,526 genes for G4, Novaseq X and Novaseq6000 and between 14,446 and 14,536 for Aviti, across all PCR cycles. The lower number of genes did not correspond to an elevated rate of PCR duplicates (Fig 9) but rather directly corresponded to the lower percentage of usable reads caused by the contamination with primer dimers (Fig 2) and with unclassified contaminants (Fig 4).

**Figure 8.**
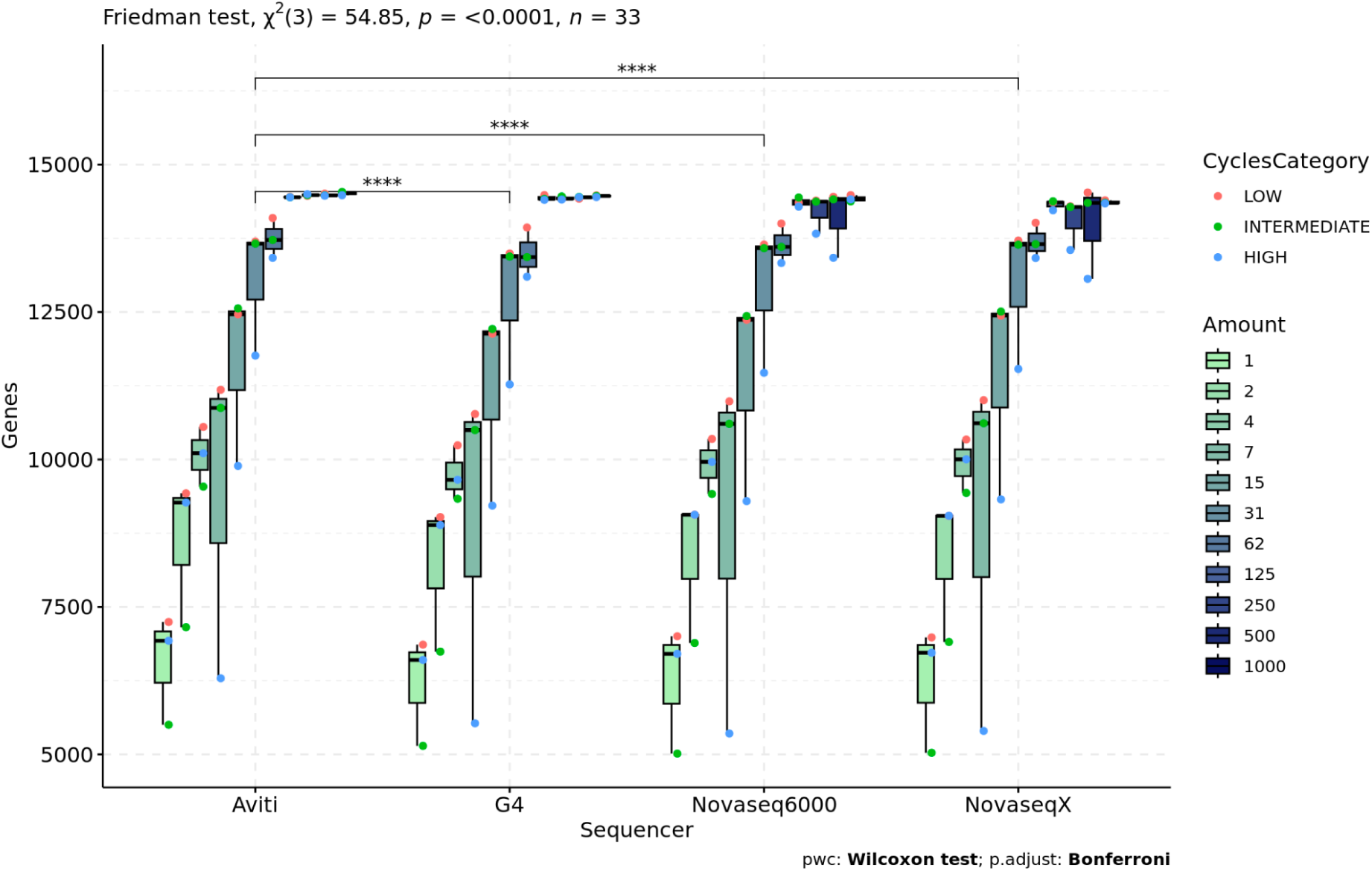
Boxplot of the number of detected genes per each input amount, PCR cycle category and sequencer. The colour of the box indicates the input amount and the colour of the points indicate the PCR cycle category. For the exact numbers refer to Supplementary Table 1. The difference in the distribution of the number of detected genes between sequencers was measured with the Friedman test. The interaction between the input amount and the number of PCR cycles was used as a blocking variable. It was followed by a paired Wilcoxon signed-rank test for pairwise comparisons between sequencers and the p-value was adjusted with the Bonferroni multiple testing correction method. The results show a significant difference in the number of detected genes between AVITI and the rest of the sequencers.

**Figure 9.**
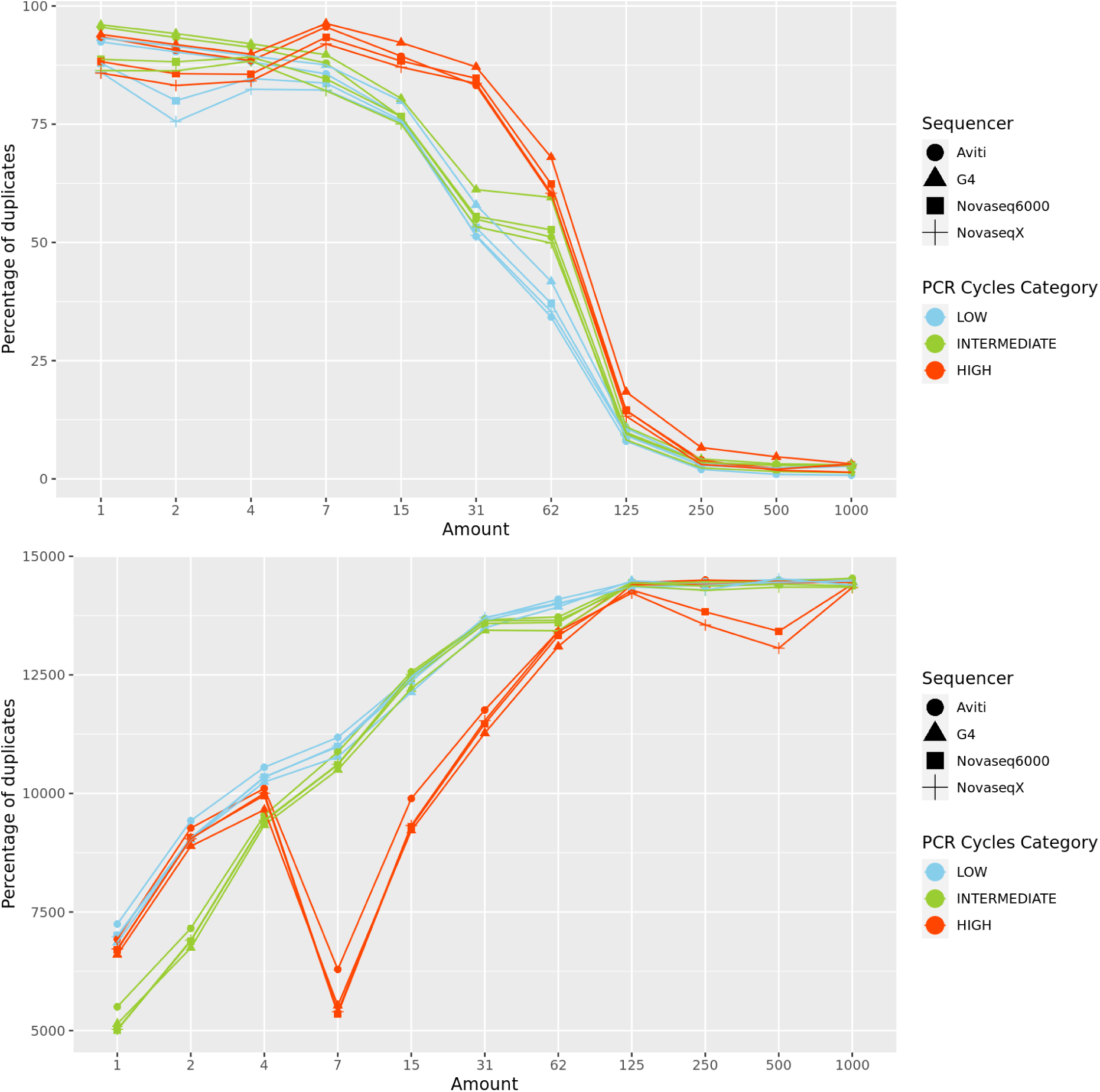
**Top**: Percentage of duplicates per sample calculated as the ratio of the mapped raw reads to the mapped deduplicated reads. The colour indicates the PCR cycle category and the shape indicates the sequencer. The data is plotted per input amount. **Bottom**: Number of detected genes for each input amount. The colour (PCR cycles category) and the shape (Sequencer) match the legend on the top plot. The number of detected genes increases with input amount. The highest PCR cycle category had the highest percentage of PCR duplicates and the lowest number of detected genes for input amounts between 7 ng and 62 ng. For input amounts above 125 ng, the percentage of duplicates was negligible for all PCR cycle categories and there was no increase in the number of detected genes.

There was a significant difference between the sequencers within the different amounts and PCR cycle categories (Friedman test, χ^2^ (3) = 55.47, p < 0.0001, n = 33, Kendall W = 0.56). Pairwise comparisons on ordered data using the paired Wilcoxon signed-rank test showed AVITI to be significantly different from the other sequences (Fig 8). The median difference in the number of genes detected by AVITI compared to G4, NovaSeq X and NovaSeq 6000 was 291, 176 and 146, respectively. However, due to the results being based on a subsampled number of reads and small number of samples, we don’t consider AVITI’s results to be biologically relevant. Additionally, in the negative control samples sequenced with AVITI, more reads remained after deduplication mapping to the human genome than for the other sequencers, which might indicate a source of contamination that impacted all samples sequenced with that platform. There was high congruency in the genes detected from each of the sequencers for the 4 highest input amounts; for the samples of 125 ng to 1,000 ng input amount, amplified with the lowest number of PCR cycles, 88% of all the genes detected were shared by at least 3 sequencers and 83% were shared by all 4 sequencers (Fig 10).

**Figure 10.**
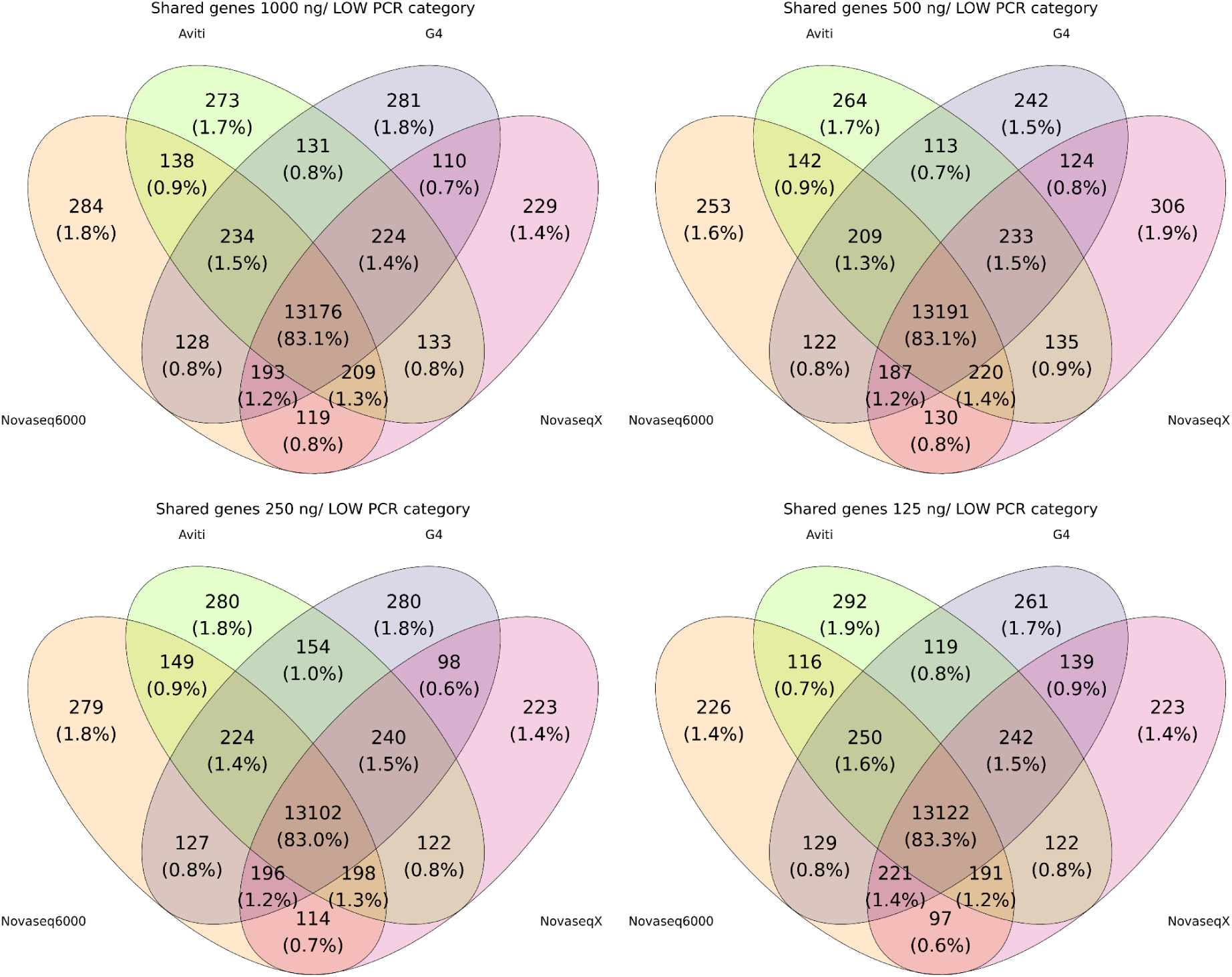
The number (and percentage) of detected genes from four highest input amounts from the lowest PCR category shared by the four sequencers. All four sequencers shared 83% of all the detected genes and 88% of the genes were detected by at least 3 sequencers.

### Gene counts are distorted by increased rate of PCR duplicates and do not differ between sequencers

Quantification of gene expression was performed with featureCounts from the Rsubread bioconductor R package, counting the number of reads mapping to each gene and producing fractional counts for multi-mapping reads. We contrasted the counts obtained for different input amounts, the PCR cycle categories and evaluated the correlation in counts between the four sequencers. We also compared the counts obtained before and after deduplication to investigate the importance of the use of UMIs in RNA-seq experiments.

We find that the combination of input amount and PCR cycle categories had an influence on the obtained counts from the unique, deduplicated reads. Low input amounts of below 31 ng showed lower counts in comparison to high input amounts of above 62 ng but with inconsistent results between the different PCR cycle categories (Fig 11). Samples of input amount between 7 ng and 31 ng amplified with the highest PCR cycle category rendered lower counts across all genes (Fig 12) and captured between 2-5% less reads in the top 20 expressed genes (Fig 13). The effect of PCR cycle categories on the counts obtained from low input amounts and the replication of that pattern across all 4 sequencers is best visualised on the UMAP (Fig 14), where the samples cluster by sequencer and separately for each PCR cycle category. For input amounts above 125 ng, the points cluster together in the centre of the plot. The counts from high input amounts were highly correspondent with a perfectly linear correlation observed between the different PCR cycle categories (Pearson’s correlation, R = 1, p-value < 0.05), except for samples of 250 ng and 500 ng (Fig 12). Samples of 250 ng and 500 ng from NovaSeq X and NovaSeq 6000 showed a deviation in counts between the High and the Low PCR cycles categories, elucidated by the low proportion of mapped reads for those two samples.

**Figure 11.**
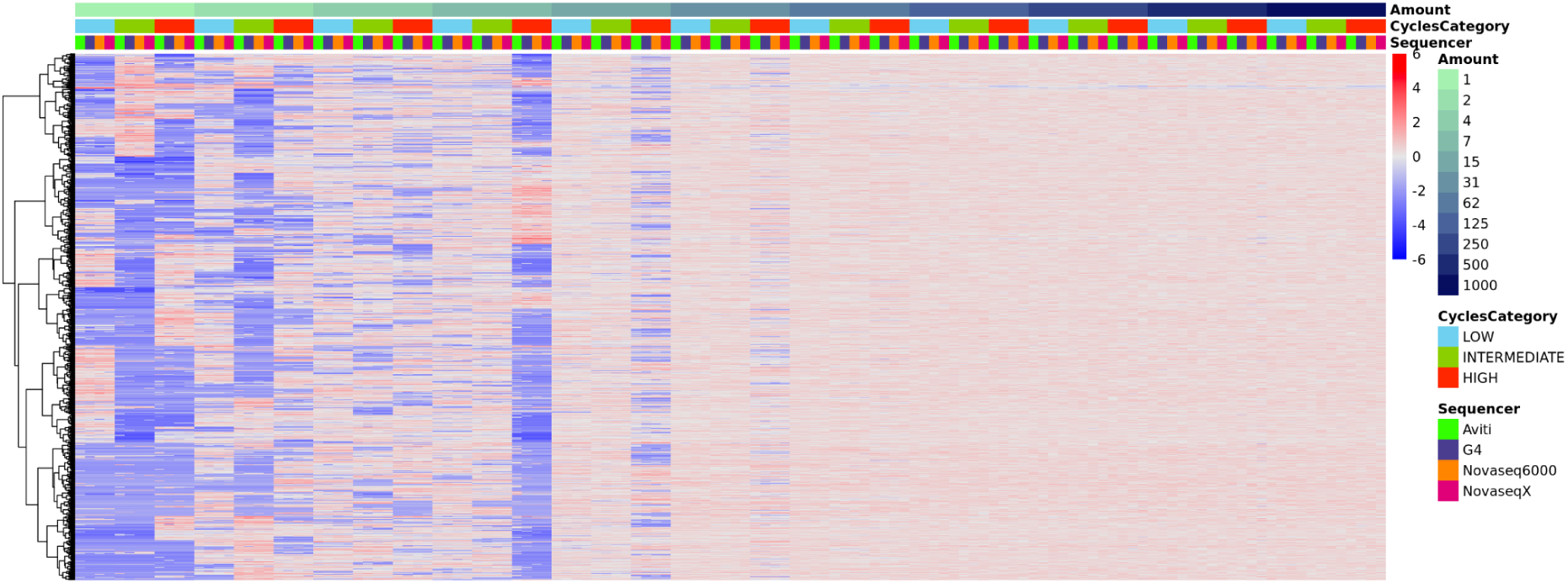
The normalised, log2 transformed counts scaled by row for the top 1000 genes with the highest variance between samples. The columns are ordered by input amount, sequencer and PCR cycles category.

**Figure 12.**
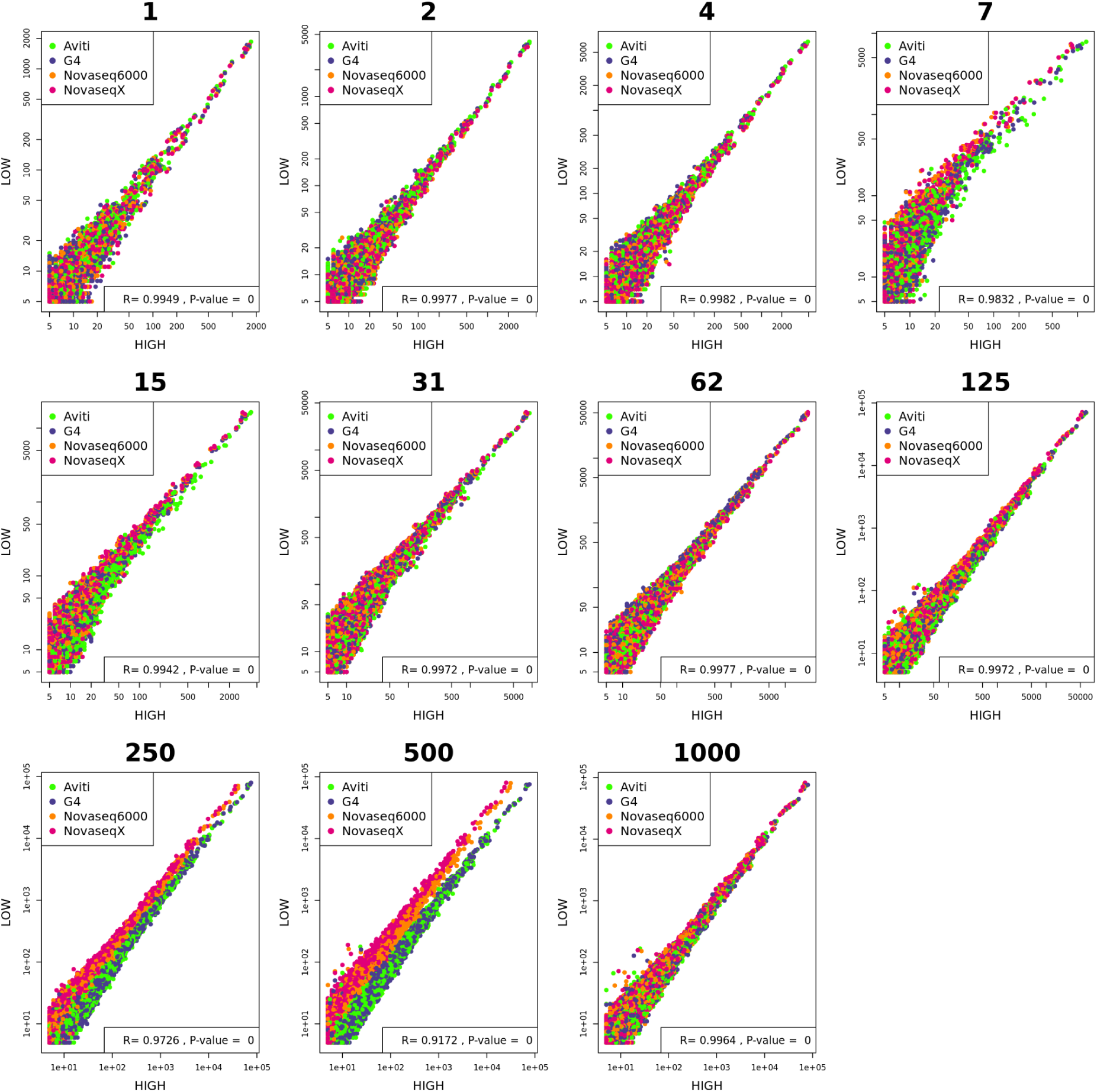
Correlation between gene counts obtained for high and low PCR cycle categories for each input amount (with an added background expression of 5). The points are coloured by the sequencer and the x and y axes are log-transformed. Pearson’s correlation coefficient (R), which determines the strength of a linear correlation (ranging from 0 to 1), is indicated, together with the p-value, at the bottom of the plot.

**Figure 13.**
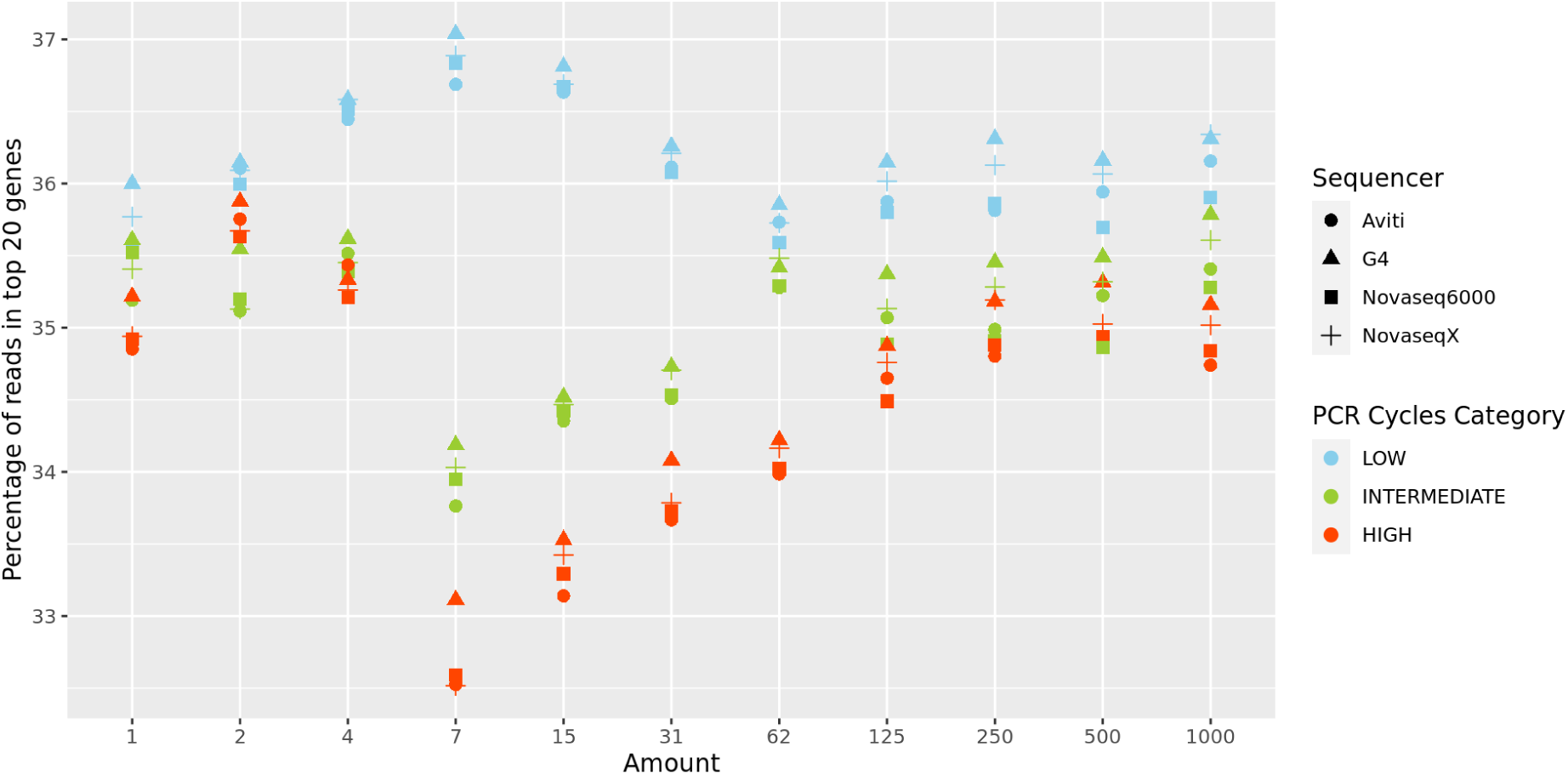
The percentage of total counts captured by the top 20 most highly expressed genes. The points are coloured by the PCR Cycles Category and the shape indicates the sequencer. High PCR cycles category captures a lower percentage of total counts in the top 20 highly expressed genes than the low PCR cycles category, most clearly for input amounts between 7 ng and 62 ng.

**Figure 14.**
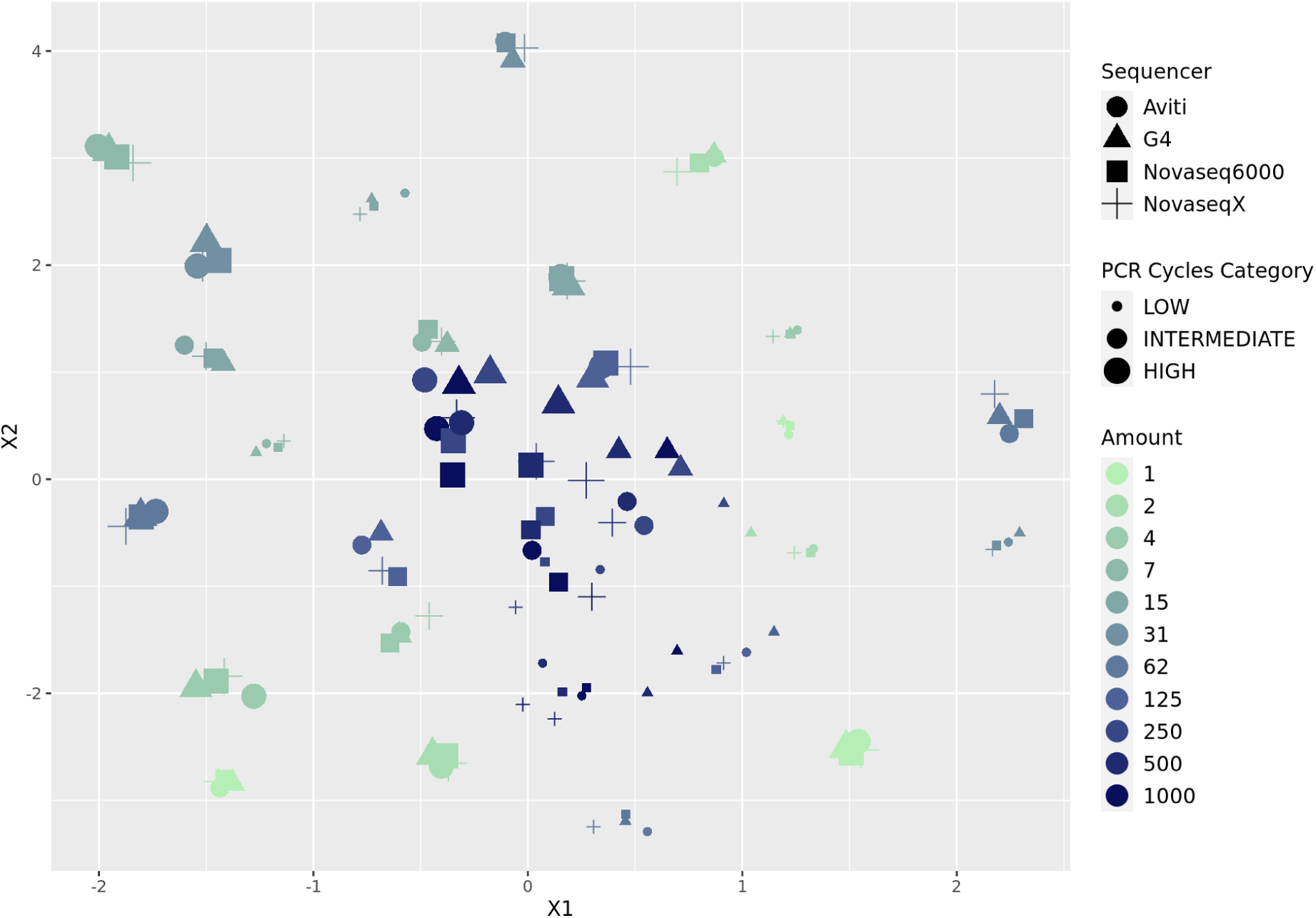
Uniform manifold approximation (UMAP) based on gene expression (normalised and log transformed gene counts obtained from featureCounts function from Rsubread R package). The points are coloured by the input amount, the size indicates the PCR cycles category and the shape indicates the sequencer. The plot shows that the PCR cycle category plays a huge role in detected counts for low input amounts (below 31 ng). For input amounts above 62 ng the effects of the PCR cycle category are not as clearly pronounced.

UMIs are widely used in the field of RNA-seq to differentiate biological copies from PCR duplicate reads (Fu et al., 2018). We used our dataset to investigate whether the use of UMIs alters the counts and if that is dependent on the input amount and PCR cycles. We used the median correlation between the technical replicates (samples sequenced with the different sequencers) to address the question (Fig 15). We find that the median correlation between sequencers increases with the increasing input amount for both the raw and the unique, deduplicated reads. Below the input of 125 ng, the correlation between the sequencers improves after deduplication of the data but for input amounts above 250 ng, there is no significant effect. Within High and Intermediate PCR cycle categories there were input amounts below 7 ng for which the correlation between the four sequencers actually decreased after deduplication, showing the inconsistency and bias in the obtained expression profile when low input amounts are used and the parameters are not optimally adjusted.

**Figure 15.**
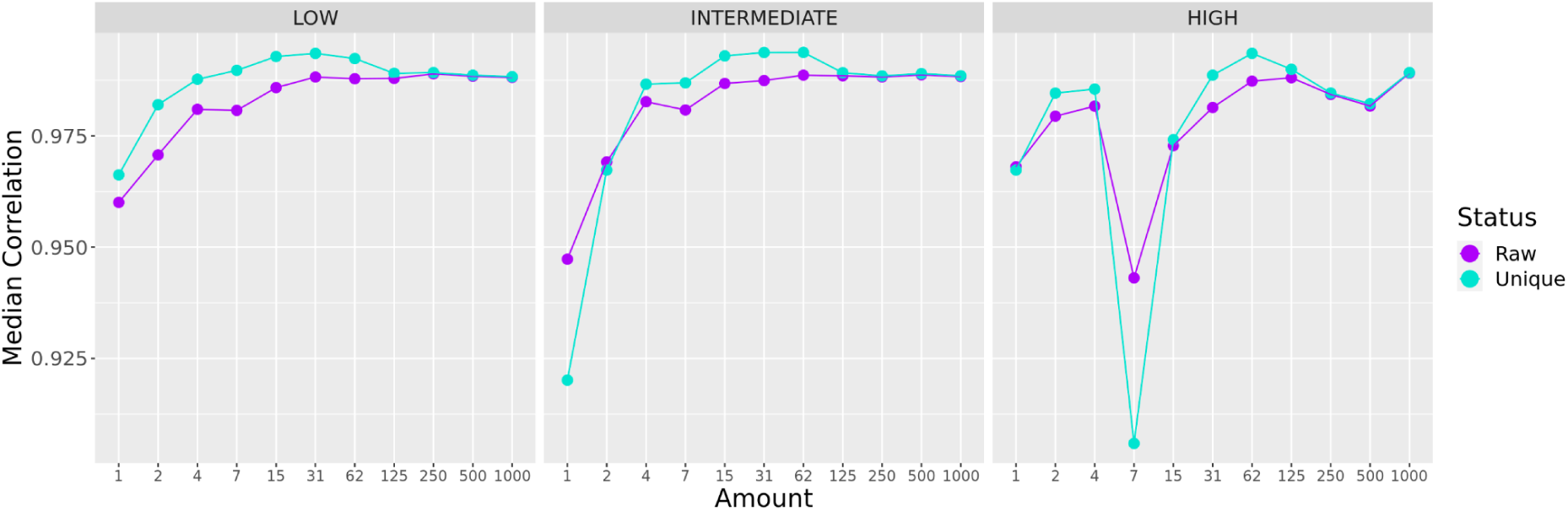
Consistency between technical replicates: median Spearman correlation coefficient from correlation of gene counts obtained from the different sequencers. The plots are limited to 1560 genes shared between all input amounts, all PCR cycle categories and all sequencers. The points are coloured by the status of the reads, raw = raw reads including duplicates, unique = unique, deduplicated reads.

The bias in the expression profiles obtained from the deduplicated reads can also be seen in the GC content of the unique, mapped reads. A deviation from the GC content distribution obtained at 1,000 ng, which follows the expected normal distribution obtained from human samples, was observed for input amounts between 1 ng and 7 ng for all PCR cycle categories and for the highest cycle category in 15 ng and 31 ng input (Fig 16).

**Fig 16.**
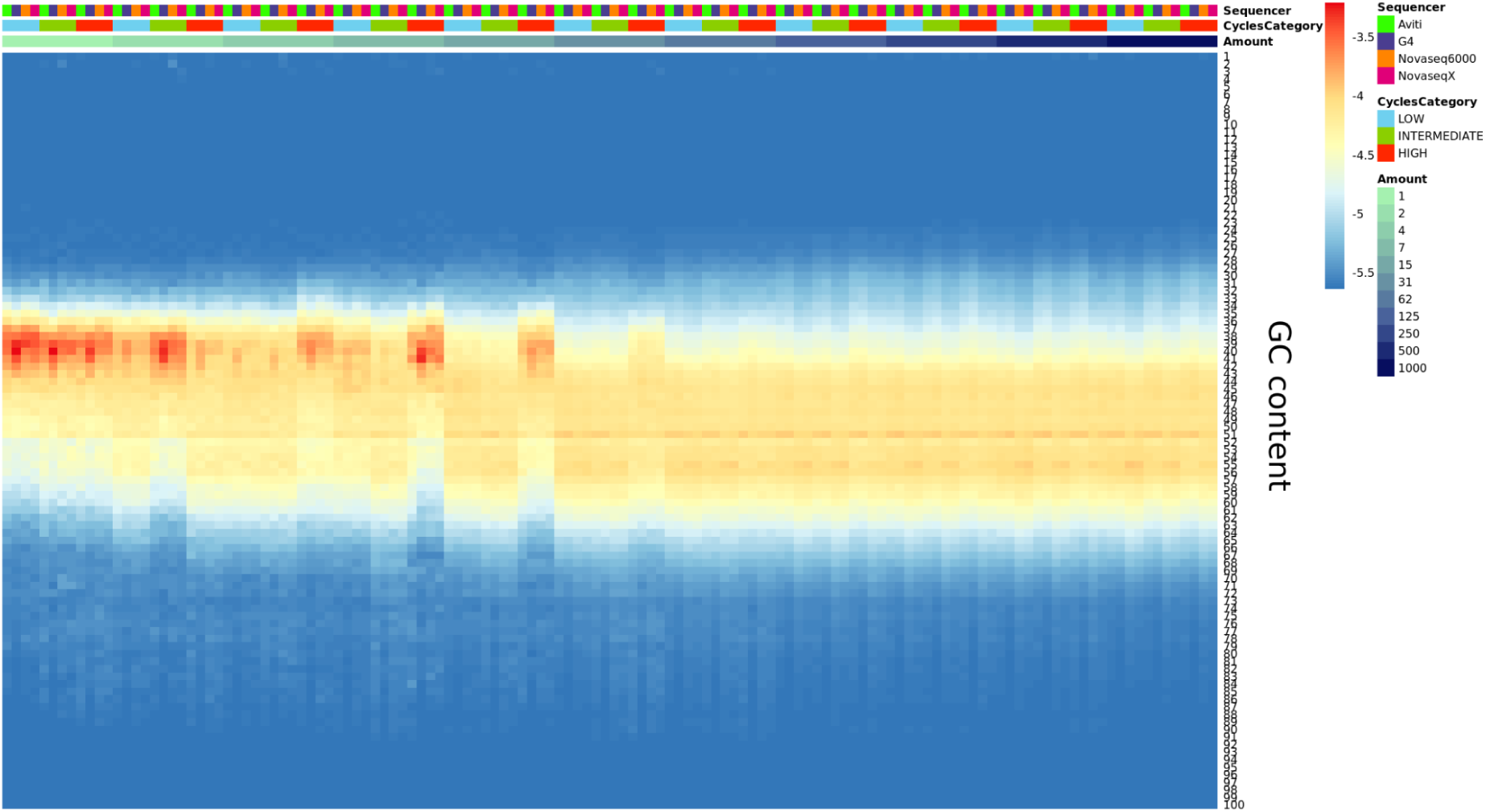
The distribution of GC content measured with picard tools across all mapped, unique reads, displayed across the different sequencers, input amounts and PCR cycle categories. The GC content is displayed on a log2 scale. Deviation from the expected GC content is observed for the low input amounts and especially for the high PCR cycle category.

## Discussion

RNA sequencing library preparation is highly sample dependent and protocol specific. One of the most important challenges in using the RNA-seq technology is how to choose an optimal set of parameters for library and sample preparation and whether variation within the recommended set of parameters heavily impacts the amount of information that can be extracted from the data after sequencing.

In this work, we focused on one of the factors falsely increasing data homozygosity and distorting transcript counts - the rate of PCR duplicates. PCR duplicates are spurious reads produced during sample amplification performed for generating enough material for sequencing (Parekh et al., 2016). We investigated whether the input amount and the number of PCR cycles, two parameters determined at the library preparation step, correlated with the production of those artefactual, unusable reads. We used the UMIs for identification and removal of PCR duplicates, additionally assessing the efficiency of one of the most widespread methods for duplicate read removal (Fu et al., 2018; You et al., 2021). We compared the number of detected genes and the gene counts obtained for each set of parameters to understand the difference in the conclusions obtained. One additional unique aspect in our study is the use of four recent short-read sequencers available on the market: NovaSeq 6000 and NovaSeq X from Illumina, AVITI from Element Biosciences and G4 from Singular Genomics. Sequencing with the two latter most novel sequencers allowed us to address the question on how the conversion kits for library preparation impact the rate of PCR duplicates and how the quality of the data compares to the widely used Illumina sequencing chemistry.

In our study, we used the NEBNext Ultra II Directional RNA Library Prep Kit protocol from Illumina for library preparation, which requires between 10 ng and 1,000 ng of input material. We find that for input amounts above 125 ng the rate of PCR duplicates is negligible and varying the number of PCR cycles applied for amplification doesn’t have an effect. We find less than 7% of the reads being identified as duplicates based on UMI and alignment coordinate. Those discarded reads did not alter the gene expression profile of the samples in a way that would impact downstream analysis.

However, we find a strong impact of the combination of the input amount and the number of PCR cycles on the rate of PCR duplicates when the starting material is below 125 ng. We observe that for the samples of 7 ng to 62 ng, the input amount is strongly negatively correlated with the proportion of PCR duplicates and thus positively correlated with the proportion of recovered, usable reads and the number of detected genes. For those samples we also observe a strong decrease in data quality when the highest number of PCR cycles is applied - we detect a much higher loss of detected genes and a deviation from the estimated gene expression in comparison to the lower value of PCR cycles. Variation in gene expression caused by unevenness of coverage introduced by amplification has already been observed in previous studies (Kozarewa et al., 2009; Ozsolak & Milos, 2011).

We find that UMI deduplication is important and effective for reliable removal of the high proportion of PCR duplicates for those samples with low input amounts, without removal of valuable biological information. For input amounts below 125 ng, between 34% and 96% of reads were discarded via deduplication. Removal of spurious reads resulted in more comparable gene expression obtained from the different PCR cycles. The highest rate of PCR duplicates and also the highest impact of deduplication was observed for input amounts below 7 ng, confirming the recommendations of the protocol. Below 13% of the reads were estimated to be productive.

We conclude that even though the protocol suggests 10 ng as the minimum input amount, we observe a huge variation in the quality of the data obtained below 125 ng with a lot more noise that needs to be corrected with the Unique Molecular Identifiers. Our results clearly demonstrate that a choice of one of the lower input amounts (15 ng) in combination with the highest number of PCR cycles can lead to a loss of even 35% of genes that could possibly be detected and can cause a surge in the rate of PCR duplicates creating noise or interference that hinders the accurate identification and measurement of the lowly to moderately expressed transcripts that in turn could be related to specific lowly expressed metabolic functions (Sheng et al., 2017).

We find a close correspondence in the results between the different sequencers in the obtained results, with minor variation. We observe the same patterns of the effect of starting material and the number of PCR cycles on PCR duplicates, similar number of detected genes and a comparable gene expression. For input amounts below 15 ng we observe a higher rate of PCR duplicates in data from AVITI and G4, suggesting an impact of the conversion kits and an additional number of PCR cycles, and so stressing higher importance of the use of UMIs in deduplicating the data. However, for input amounts below 15 ng we observe a higher proportion of reads filtered due to length (< 18bp) in the data from both of the Illumina sequencers. For the G4 sequencer we observe an elevated sequencing error rate measured through the number of mismatches in the mapped raw reads. However, we do not see any impact of that on downstream results including the mapping rate, number of detected genes or gene expression profiles. We also find a significantly but marginally higher number of genes detected in data from AVITI when compared to the other sequencers. For some input amounts, both high and low, we do see a slightly higher mapping rate and lower rate of PCR duplicates in the data from AVITI. However, due to only one sample representing each set of parameters and the data being subsampled, we are unable to comment on the significance of this finding. Additionally, the negative control sample sequenced with Aviti had an elevated number of reads remaining after deduplication that mapped to the human genome than the other sequencers, which might indicate a source of contamination that impacted all samples sequenced with that platform.

One noticeable difference between the sequencers was the presence of contamination by adapter primer dimers. Two samples from NovaSeq 6000 and NovaSeq X with input amounts of 250 ng and 500 ng did not exactly match the high quality results from the rest of the samples due to a higher proportion of adapter primer dimers. The effect was most prominently visible when the samples were amplified using the highest PCR cycle category. During the conversion of the Illumina library to a library suitable for sequencing on AVITI and G4, the primer dimers were cleaned up. These samples serve as an example that with an increase in the number of PCR cycles for amplification, the rate of adapter primer dimerization also increases (Bhargava et al., 2014). To avoid wasting sequencing efforts and production of low quality data, size selection in library preparation could be applied to filter out the contaminating primer dimers. However, one has to note that size selection itself can introduce transcript length bias also resulting in complications in downstream analyses (Ozsolak & Milos, 2011).

Low sample input amounts, which are often seen in limiting samples processed in many core facilities, utilise amplification methods during library preparation and result in high rate of PCR duplicates. The rate of PCR duplicates also increases if Illumina native libraries of low input amounts have to be converted for sequencing on other sequencers, nowadays more frequently available. Our research shows that PCR duplicate removal through the use of UMIs results in data with less noise and bias, thereby increasing FDR and power of downstream analyses. Removal of the artefactual reads also results in comparable data across all four short-read sequencers.

## Acknowledgements

● Lutz Froenicke, DNA Technologies and Expression Analysis Core UC Davis Genome Center Element AVITI sequencing

● Laura Neff, Joel Wirz, Hai Bui, FGCZ Genomics NGS Team for the NovaSeq 6000 and NovaSeq X sequencing

● Darius Fugere and Lauren Moller, Singular Genomics for technical support

● Robert Steen, member of the Genomics Research Group of the Association of Biomolecular Resource Facilities (ABRF) for the early discussions.

● Active members of the Genomics Research Group of the Association of Biomolecular Resource Facilities (ABRF)

## Notes

### Competing Interest Statement

The authors have declared no competing interest.

