## Supplementary material for "The impact of PCR duplication on RNAseq data generated using NovaSeq 6000, NovaSeq X, AVITI and G4 sequencers": Analysis.Rmd

````
---
title:  "PCR Duplication rates"
author: "Natalia Zajac/ Hubert Rehrhauer"
output:
  html_document: 
    highlight: pygments
    theme: sand
    code_folding: hide
    toc: yes
    toc_depth: 4
editor_options: 
  chunk_output_type: inline
---

```{r setup countQC, include=FALSE}
knitr::opts_chunk$set(warning = FALSE, message = FALSE) 
library(ggplot2)
library(plotly)
library(ggrepel)
library(ggpubr)
library(grid)
library(gridExtra)
library(ezRun)
library(ggtreeExtra)
library(ggtree)
library(dplyr)
library(tidyverse)
library(cowplot)
library(scales)
library(DESeq2)
library(EnhancedVolcano)
library(edgeR)
library(UpSetR)
library(pheatmap)
library(ggvenn)
library(umap)
knitr::opts_chunk$set(echo = TRUE, warning = FALSE, message = FALSE, knitr.table.format = "html")
#BPPARAM <- MulticoreParam(workers = 4)
#register(BPPARAM)
```


```{r, echo=FALSE}
#load the data

baseDir <- "/srv/GT/analysis/zajacn/p2220"
dedupFolders <- c(Novaseq6000="o31444/o31444_NovaSeq_NOV1703_subsampled_dedup",
                         Aviti="o31444_Aviti/Aviti_p2220_o31444_subsampled_dedup",
                         G4="o31444_G4/G4_p2220_o31444_subsampledv2_dedup",
                         NovaseqX="o31444_NovaseqX_100bp/dedup")
geneAnno <- ezRead.table("/srv/GT/reference/Homo_sapiens/GENCODE/GRCh38.p13/Annotation/Release_42-2023-01-30/Genes/genes_annotation_byGene.txt")
protcodingAnno <- geneAnno[geneAnno$type == "protein_coding",]


tablex = data.frame()
for (nm in names(dedupFolders)){
  file = list.files(path = file.path(baseDir, dedupFolders[nm]), 
                    pattern = "flagstat.primary.summary.table.txt", full.names = TRUE)
  table <- ezRead.table(file)
  table$removed <- table$ReadsIn - table$ReadsOut
  table$Proportion_of_reads_removed <- (table$removed/table$ReadsIn)*100
  table <- table %>% rownames_to_column() %>% separate(rowname, c("Amount", "cycles"), "_")
  table = table %>% mutate(Sequencer = nm)
  table$cycles <- as.integer(table$cycles)
  table$Prededuplication <- (table$ReadsIn/table$ReadNumber)*100
  table$Postdeduplication <- (table$ReadsOut/table$ReadNumber)*100
  tablex = rbind(tablex,table)
}

sampleInfo <- tablex[,c(1,2,8)] %>% 
  mutate(Id = paste0(Amount, "_", cycles, "_", Sequencer)) %>% 
  mutate(amountFac= if_else(Amount == "0", "NC", Amount))
sampleInfo$amountFac <- sampleInfo$amountFac %>%
  factor(levels= c("NC", "1", "2", "4", "7", "15", "31", "62", "125", "250", "500", "1000"))
sampleInfo$`Cycles Category` = ""

for (i in unique(sampleInfo$Amount)){
  sampleInfo[(sampleInfo$Amount == i) & (sampleInfo$cycles == min(as.integer(sampleInfo[sampleInfo$Amount == i,]$cycles))),"Cycles Category"] <- "LOW"
  sampleInfo[(sampleInfo$Amount == i) & (sampleInfo$cycles == max(as.integer(sampleInfo[sampleInfo$Amount == i,]$cycles))),"Cycles Category"] <- "HIGH"
  sampleInfo[(sampleInfo$Amount == i) & (sampleInfo$`Cycles Category` == ""),"Cycles Category"] <- "INTERMEDIATE"
}
sampleInfo$`Cycles Category` <- factor(sampleInfo$`Cycles Category`, levels=c("LOW", "INTERMEDIATE", "HIGH"))
sampleInfo$Sample = paste0(sampleInfo$Amount, "_", sampleInfo$cycles)
tablex = merge(tablex, sampleInfo, by = c("Amount", "cycles", "Sequencer"))


rawCounts <- NULL
dedupCounts <- NULL
dataInfo <- NULL
nm <- names(dedupFolders[1])
for (nm in names(dedupFolders)){
  file = list.files(path = file.path(baseDir, dedupFolders[nm]), 
                    pattern = "featurecounts.pre.post.combined.txt", full.names = TRUE)
  combinedMatrix <- ezRead.table(file, quote="\"") ## why is that matrix quoted???
  x <- combinedMatrix[ , grep("_pre", colnames(combinedMatrix))]
  colnames(x) <- sub("pre", nm, colnames(x))
  if (is.null(rawCounts)){
    rawCounts <- x
  } else {
    rawCounts <- cbind(rawCounts, x)
  }
  x <- combinedMatrix[ , grep("_post", colnames(combinedMatrix))]
  colnames(x) <- sub("post", nm, colnames(x))
  if (is.null(dedupCounts)){
    dataInfo <- ezFrame(sampleInfo)
    dedupCounts <- x
  } else {
    dedupCounts <- cbind(dedupCounts, x)
    dataInfo <- rbind(dataInfo, ezFrame(sampleInfo))
  }
}
dataInfo = dataInfo[!duplicated(dataInfo),]
stopifnot(setequal(dataInfo$Id, colnames(dedupCounts)))
stopifnot(rownames(protcodingAnno) %in% rownames(dedupCounts))

dedupCounts <- as.matrix(dedupCounts[ rownames(protcodingAnno), dataInfo$Id])
rawCounts <- as.matrix(rawCounts[ rownames(protcodingAnno), dataInfo$Id])

isAllPresent <- apply(dedupCounts[,colnames(dedupCounts)[!grepl("^0_", colnames(dedupCounts))]] > 0, 1, all)
sum(isAllPresent)

tablex = merge(tablex, as.data.frame(colSums(dedupCounts > 0)), by.x = "Id", by.y = "row.names") %>% dplyr::rename("Detected_genes" = 'colSums(dedupCounts > 0)')
```


```{r, echo=FALSE}
starFolders <- 
  c(Novaseq6000="/srv/gstore/projects/p2220/o31444_STAR_markdups_Novaseq_subsampled_2023-08-24--10-13-19",
                         Aviti="/srv/gstore/projects/p2220/STAR_markdups_Aviti_subsampled_2023-08-24--10-09-41",
                         G4="/srv/gstore/projects/p2220/STAR_markdups_G4_subsampled_2023-08-24--10-16-09",
                         NovaseqX="/srv/gstore/projects/p2220/o31444_STAR_cutto100bp_2023-12-07--17-39-06")


readCountList <- list()
for (seqr in names(starFolders)){
  starDsFile <- file.path(starFolders[seqr], "dataset.tsv")
  starDs <- ezRead.table(starDsFile)
  rownames(starDs) <- sub("_S.*", "", rownames(starDs))
  stopifnot(setequal(sampleInfo$Sample, rownames(starDs)))
  sm <- sampleInfo$Sample[1]
  resList <- list()
  for (sm in sampleInfo$Sample){
    logFile <- file.path("/srv/gstore/projects", starDs[sm, "PreprocessingLog [File]"])
    logLines <- readLines(logFile, warn = FALSE)
    jsonStart <- grep("^\\{", logLines)
    x <- jsonlite::fromJSON(paste(logLines[jsonStart:length(logLines)], collapse="\n"))
    nReads <- x$summary$before_filtering$total_reads
    xStat <- c(Sequencer=seqr, total_reads=nReads, x$filtering_result)
    resList[[sm]] <- xStat
  }
  readCountList[[seqr]] <- rbindlist(resList, idcol="Sample") %>% data.frame()
}

readCounts <- rbindlist(readCountList) %>% data.frame()
readCounts$all_low_quality_reads <- readCounts$low_quality_reads + readCounts$too_many_N_reads
readCounts$low_quality_reads <- NULL
readCounts$too_many_N_reads <- NULL
readCounts$too_long_reads <- NULL

xx = merge(tablex, readCounts, by = c("Sample", "Sequencer"))
xx = merge(xx, round(colSums(dedupCounts)), by.x = "Id", by.y = "row.names") %>% dplyr::rename("uniqueMappedCounted" = "y")
xx$uniqueMappedNotCounted <- round(xx$ReadsOut - xx$uniqueMappedCounted)
xx$dupsMapped <- xx$ReadsIn - xx$ReadsOut
xx$unMapped <- xx$passed_filter_reads - xx$ReadsIn
xx$lowQuality <- xx$all_low_quality_reads
xx$tooShort <- xx$too_short_reads
xLong <- reshape2::melt(xx[,c("Sample", "Sequencer", "amountFac", "Cycles Category", "uniqueMappedCounted",  "uniqueMappedNotCounted", "dupsMapped", "unMapped", "lowQuality", "tooShort")])
xLong$Sample <- factor(xLong$Sample, levels=unique(xx$Sample))
xLong$PCR <- as.character(xLong$`Cycles Category`)

```

## Figure 2
```{r , fig.width = 19, fig.height=13, echo=FALSE}
xLong$CyclesCategory = str_replace(xLong$`Cycles Category`, "INTERMEDIATE", "MID")
xLong$CyclesCategory = factor(xLong$CyclesCategory, levels = c("LOW", "MID", "HIGH"))
ggplot(xLong, aes(x=CyclesCategory, y=value, fill=variable)) + 
  geom_bar(stat="identity", position="fill") + 
  facet_grid(rows = vars(Sequencer), cols = vars(amountFac)) +
  scale_fill_discrete("Category",labels=c('unique reads mapped and counted', 'unique reads mapped and not counted', "duplicate reads mapped", "unmapped reads", "low quality reads", "reads < 18bp")) +
  labs(y = "Proportion", x = "PCR Cycles Category") + theme(axis.text.x = element_text(size = 12), strip.text = element_text(size = 15), legend.text = element_text(size = 18), legend.title = element_text(size = 18), legend.position = "top")
```

##Figure 3
```{r , fig.width = 12, fig.height=5, echo=FALSE}
#Run qualimap
resBaseDir <- "/srv/GT/analysis/p2220/PCR-duplicates-study/qualimap_NovaseqX100bp/"
# if (!file.exists(resBaseDir)){
#   dir.create(resBaseDir, recursive = TRUE)
# }
# 
# for (seqr in names(starFolders)){
#   starDsFile <- file.path(starFolders[seqr], "dataset.tsv")
#   starDs <- ezRead.table(starDsFile)
#   rownames(starDs) <- sub("_S.*", "", rownames(starDs))
#   stopifnot(setequal(sampleInfo$Sample, rownames(starDs)))
#   for (sm in sampleInfo$Sample){
#     bamFile <- file.path("/srv/gstore/projects", starDs[sm, "BAM [File]"])
#     resDir <- paste0(resBaseDir, "/", sm, "_", seqr)
#     message(resDir)
#     cmd <- paste("unset DISPLAY; /usr/local/ngseq/packages/QC/Qualimap/2.2.1/qualimap bamqc -bam", bamFile, "--java-mem-size=4G", "-c -nw 400 -hm 3 -nt 8", "-outdir", resDir)
#     ezSystem(cmd)
#     }
# }

mismatchRate <- ezMatrix(NA, rows=unique(sampleInfo$Sample), cols=names(starFolders))
for (seqr in names(starFolders)){
    for (sm in sampleInfo$Sample){
        resDir <- paste0(resBaseDir, "/", sm, "_", seqr)
        genomeResFile <- paste0(resDir, "/genome_results.txt")
        x <- readLines(genomeResFile)
        errorRate <- grep("general error rate = ", x, value=TRUE) %>% sub(".*= ", "", x=.) %>% as.double()
        mismatchRate[sm, seqr] <- errorRate
    }
}
rownames(mismatchRate) <- unique(paste0(sampleInfo$amountFac, "_", sampleInfo$`Cycles Category`))
mismatchRate = reshape2::melt(mismatchRate)
mismatchRate$Var1 = factor(mismatchRate$Var1, levels = unique(paste0(dataInfo[with(dataInfo, order(amountFac, `Cycles Category`)),]$amountFac, "_", dataInfo[with(dataInfo, order(amountFac, `Cycles Category`)),]$`Cycles Category`)))

ggplot(mismatchRate, aes(x=Var1, y=value, group=Var2, color=Var2)) +
  geom_step(position="jitter", direction="mid") +
  theme(axis.text.x = element_text(angle = 45, vjust = 1, hjust=1)) + 
  labs(x="SampleID", y = "Qualimap Mismatch Rate", color = "Sequencer") + 
  scale_color_manual(values = c("darkorange", 'chartreuse','darkslateblue', "deeppink3"))

```

## Figure 4
```{r , fig.width = 18, fig.height=10, echo=FALSE}
all_bracken = NULL
for (dir in list.dirs()[ grepl("contamination$", list.dirs()) ]){
  bracken = NULL
  for (x in list.files(dir, pattern = "txt.bracken")){
    X = read.delim(paste0(dir,"/",x)) %>% mutate(name = if_else(name == "Homo", "Homo sapiens", name))
    X$SampleID = str_split(x, "_S[1-9]+")[[1]][1]
    bracken = rbind(bracken, X)
  }
  bracken$Sequencer = str_split(dir, "/")[[1]][2]
  all_bracken = rbind(all_bracken, bracken)
}
all_bracken = all_bracken[all_bracken$fraction_total_reads > 0.001,]

all_kraken = NULL
for (dir in list.dirs()[ grepl("contamination$", list.dirs()) ]){
  X = read.delim(paste0(dir,"/kraken_read_summary.txt"), header = FALSE)
  colnames(X) = c("SampleID", "Classified", "Unclassified")
  X$Sequencer = str_split(dir, "/")[[1]][2]
  all_kraken = rbind(all_kraken, X)
}
all_kraken$SampleID = str_remove(all_kraken$SampleID, "_S[1-9]+")

all_bracken = all_bracken[,c(1,6,8,9)]
all_kraken = all_kraken[,c(1,3,4)] %>% dplyr::rename("new_est_reads" = "Unclassified") %>% mutate(name = "Unclassified")
contamination = rbind(all_bracken, all_kraken)

contamination = contamination %>% 
  mutate(Sequencer = case_when(Sequencer == "o31444_Aviti" ~ "Aviti",
                                          Sequencer == "o31444_G4" ~ "G4",
                                          Sequencer == "o31444_NovaseqX" ~ "NovaseqX",
                                          Sequencer == "o31444" ~ "Novaseq6000")) %>% 
  separate(SampleID, into = c("Amount", "cycles"), sep = "_")
contamination$name = factor(contamination$name, levels = unique(contamination[order(contamination$new_est_reads, decreasing = T),]$name))
contamination$Amount <- str_replace(as.character(contamination$Amount), "^0$", "NC")
contamination$Amount = factor(contamination$Amount , levels = c("NC", "1", "2", "4", "7", "15", "31", "62", "125", "250", "500", "1000"))
contamination = contamination %>% group_by(Amount, Sequencer, name) %>% summarise(mean = mean(new_est_reads))

ggplot(contamination, aes(Amount, mean, fill = name)) + 
  geom_bar(position = "fill", stat = "identity") + 
  facet_wrap(~Sequencer) + ylab("Total") + 
  theme(title = element_text(size = 25), 
        axis.text.x = element_text(size = 12), 
        axis.text.y = element_text(size = 15), 
        legend.text = element_text(size = 13), 
        strip.text = element_text(size=25)) + 
  labs(fill="Read Classification") +
  scale_fill_manual(values = sample(ezRun::brewPalette(n=150),85)) + 
  guides(fill = guide_legend(ncol = 2))
```

### Figure 5
```{r, fig.width = 17, fig.height = 6, echo=FALSE}

ggplot(tablex, aes(x = amountFac, y = ReadNumber, color = Sequencer, group = Sequencer)) + 
  facet_wrap(~`Cycles Category`) + 
  geom_point(size = 3.5) + 
  scale_color_manual(values=c('chartreuse','darkslateblue', "darkorange","deeppink3")) + 
  geom_line() + 
  xlab("Amount") + 
  theme(axis.text.x = element_text(angle= 45, size = 15, hjust= 1), text = element_text(size = 17), axis.text.y = element_text(size = 15), legend.text = element_text(size = 15)) +
  ylim(0,2000000) +
  ylab("Read Number")
```


## Figure 6
```{r, fig.width = 17, fig.height = 6, echo=FALSE}
df1 = melt(tablex[,c("amountFac", "Cycles Category", "Sequencer", "Prededuplication", "Postdeduplication")],id.vars=c("amountFac","Cycles Category","Sequencer"))
df1 <- df1 %>% dplyr::rename("Status" = "variable")

F2 = ggplot(df1, 
       aes(x = amountFac, y = value, col = Status, shape = Sequencer ,group = interaction(Status, Sequencer))) + 
  xlab("Amount") + 
  ylab("Reads mapped %") + 
  facet_wrap(~`Cycles Category`) + 
  geom_point(size = 3) + 
  geom_line() + 
  theme(axis.text.x = element_text(size = 10), text = element_text(size = 17)) +
  scale_color_manual(values=c('Purple','Turquoise'), labels=c('Raw', 'Unique')) + 
  guides(shape = guide_legend(override.aes = list(size = 5)), color = guide_legend(override.aes = list(size = 5)))
F2
```

### Figure 7
```{r , fig.width = 12, fig.height=8, echo=FALSE}
if(!exists("dataset")){
  dataset <- readRDS("/srv/gstore/projects/p2220/RNABamStats_2023-12-09--17-33-58/RNA_BAM_Statistics/dataset.rds")
}
if(!exists("param")){
  param <- readRDS("/srv/gstore/projects/p2220/RNABamStats_2023-12-09--17-33-58/RNA_BAM_Statistics/param.rds")
}
if(!exists("resultList")){
  resultList <- readRDS("/srv/gstore/projects/p2220/RNABamStats_2023-12-09--17-33-58/RNA_BAM_Statistics/resultList.rds")
}

conds = ezConditionsFromDataset(dataset, param=param)
samples = rownames(dataset)
sampleColors = getSampleColors(conds, samples)
bamFiles = dataset$BAM

nm="multiMatchTargetTypeCounts"
tct = getTypeCountTable(resultList, nm)
tpt = as.matrix(tct)
for (cn in colnames(tpt)){
    tpt[ ,cn] = tct[ ,cn]/ tct["total", cn] * 100
}
eliminate = c(rownames(tpt)[grepl("chr", rownames(tpt))], rownames(tpt)[grepl("IG_", rownames(tpt))], rownames(tpt)[grepl("TR_", rownames(tpt))], "total", "transcribed_processed_pseudogene", "TEC", "transcribed_unprocessed_pseudogene", "transcribed_unitary_pseudogene", "vault_RNA", "ribozyme", "snRNA", "rRNA", "miRNA", "misc_RNA", "snoRNA", "scaRNA", "sRNA", "scRNA", "Mt_tRNA")
tptUse = tpt[rownames(tpt)[!rownames(tpt) %in% eliminate],]

ggplot(melt(tptUse) %>% rownames_to_column(), aes(value, Var1, fill = Var2)) + 
    geom_bar(stat = "identity", position = "dodge", width = 0.5) +
    scale_fill_manual(values = c(hsv(1, seq(0,1,length.out = 12)), hsv(0.5, .35, seq(.45,.90,length.out = 12)))) +
    labs(x = "Percentage of mapped unique reads", y = "Category", fill="SampleID")
```

## Figure 8
```{r , fig.width = 11, fig.height=7, echo=FALSE}
library(rstatix)
library(viridis)
df = tablex[tablex$amountFac != "NC", c("amountFac", "Cycles Category", "Sample", "Sequencer", "Detected_genes")]
df$Amount = factor(df$amountFac, levels = c("1", "2", "4", "7","15","31", "62", "125", "250", "500","1000"))
df  = df %>% dplyr::rename("CyclesCategory" = "Cycles Category") %>% dplyr::rename("Genes" = 'Detected_genes')
df$CyclesCategory = factor(df$CyclesCategory, levels = c("LOW", "INTERMEDIATE", "HIGH"))
df$inter <- interaction(as.factor(df$Amount), as.factor(df$CyclesCategory))
df$Sequencer = factor(df$Sequencer, levels = c("Aviti", "G4", "Novaseq6000", "NovaseqX"))
df = df %>% arrange(Sequencer, Amount, CyclesCategory)
res = df %>% friedman_test(Genes ~ Sequencer | inter)
pwc <- df %>%
    wilcox_test(Genes ~ Sequencer, paired = TRUE, p.adjust.method = "bonferroni")
pwc <- pwc %>% add_xy_position(x = "Sequencer", dodge = 0.8)
ggboxplot(df, x = "Sequencer", y = "Genes", fill = "Amount") +
    stat_pvalue_manual(pwc, hide.ns = TRUE,tip.length = 0.02,
                       step.increase = 0.1) +
    labs(
        subtitle = get_test_label(res,  detailed = TRUE),
        caption = get_pwc_label(pwc)
    ) + scale_fill_manual(values = carto.pal(pal1 = "turquoise.pal" ,n1 = 11)) + geom_point(position=position_dodge(width=0.75),aes(group=Amount, color = CyclesCategory))  +
    grids(linetype = "dashed") + theme(legend.position = "right")

list_of_G4 = list()
list_of_NX = list()
list_of_N6 = list()
for (i in unique(df$amountFac)){
  for (x in unique(df$CyclesCategory)){
    value = df[(df$amountFac == i) & (df$CyclesCategory == x) & (df$Sequencer == "Aviti"),]$Genes - df[(df$amountFac == i) & (df$CyclesCategory == x) & (df$Sequencer == "G4"),]$Genes
    list_of_G4 = c(list_of_G4, value)
    value = df[(df$amountFac == i) & (df$CyclesCategory == x) & (df$Sequencer == "Aviti"),]$Genes - df[(df$amountFac == i) & (df$CyclesCategory == x) & (df$Sequencer == "NovaseqX"),]$Genes
    list_of_NX = c(list_of_NX, value)
    value = df[(df$amountFac == i) & (df$CyclesCategory == x) & (df$Sequencer == "Aviti"),]$Genes - df[(df$amountFac == i) & (df$CyclesCategory == x) & (df$Sequencer == "Novaseq6000"),]$Genes
    list_of_N6 = c(list_of_N6, value)
  }
}

median(unlist(list_of_G4))
median(unlist(list_of_NX))
median(unlist(list_of_N6))

```

##Figure 9
```{r , fig.width = 10, fig.height=10, echo=FALSE}

p = ggplot(tablex[tablex$Amount != "0",], aes(factor(amountFac,levels= c("1", "2", "4", "7", "15", "31", "62", "125", "250", "500", "1000")), Proportion_of_reads_removed, color=`Cycles Category`, shape=Sequencer, group = interaction(`Cycles Category`, Sequencer))) + geom_point(size = 2) + labs(x = "Amount", y = "Percentage of duplicates", color = "PCR Cycles Category") + guides(color = guide_legend(override.aes = list(size = 5)), shape = guide_legend(override.aes = list(size = 5))) + geom_line() + scale_color_manual(values = c("skyblue", "yellowgreen", "orangered")) 
p1 = ggplot(tablex[tablex$Amount != "0",], aes(factor(amountFac,levels= c("1", "2", "4", "7", "15", "31", "62", "125", "250", "500", "1000")), Detected_genes, color=`Cycles Category`, shape=Sequencer, group = interaction(`Cycles Category`, Sequencer))) + geom_point(size = 2) + labs(x = "Amount", y = "Percentage of duplicates", color = "PCR Cycles Category") + guides(color = guide_legend(override.aes = list(size = 5)), shape = guide_legend(override.aes = list(size = 5))) + geom_line() + scale_color_manual(values = c("skyblue", "yellowgreen", "orangered")) 
plot_grid(p, p1, ncol = 1) 
```

##Figure 10
```{r , fig.align = "center", fig.width = 20, fig.height=16, echo=FALSE, out.width='\\textwidth'}
dfgenes = NULL
for (i in colnames(dedupCounts)) {
        dfgenes[[i]] = rownames(dedupCounts[dedupCounts[,i] != 0,])
}

one = list("Novaseq6000" = dfgenes$'1000_6_Novaseq6000', "Aviti" = dfgenes$'1000_6_Aviti', "G4" = dfgenes$'1000_6_G4', "NovaseqX" = dfgenes$'1000_6_NovaseqX')
two = list("Novaseq6000" = dfgenes$'500_7_Novaseq6000', "Aviti" = dfgenes$'500_7_Aviti', "G4" = dfgenes$'500_7_G4', "NovaseqX" = dfgenes$'500_7_NovaseqX')
three = list("Novaseq6000" = dfgenes$'250_8_Novaseq6000', "Aviti" = dfgenes$'250_8_Aviti', "G4" = dfgenes$'250_8_G4', "NovaseqX" = dfgenes$'250_8_NovaseqX')
four = list("Novaseq6000" = dfgenes$'125_9_Novaseq6000', "Aviti" = dfgenes$'125_9_Aviti', "G4" = dfgenes$'125_9_G4', "NovaseqX" = dfgenes$'125_9_NovaseqX')

venn1 <- ggvenn(
    one,
    set_name_size = 10, text_size = 15, fill_alpha = 0.2, stroke_size = 0.5) + ggtitle("Shared genes 1000 ng/ LOW PCR category") +
  theme(plot.title = element_text(hjust = 0.5, size = 35)) + scale_fill_manual(values = c("darkorange", 'chartreuse','darkslateblue', "deeppink3"))
venn2 <- ggvenn(
    two,
    set_name_size = 10, text_size = 15, fill_alpha = 0.2, stroke_size = 0.5) + ggtitle("Shared genes 500 ng/ LOW PCR category") +
  theme(plot.title = element_text(hjust = 0.5, size = 35)) + scale_fill_manual(values = c("darkorange", 'chartreuse','darkslateblue', "deeppink3"))
venn3 <- ggvenn(
    three,
    set_name_size = 10, text_size = 15, fill_alpha = 0.2, stroke_size = 0.5) + ggtitle("Shared genes 250 ng/ LOW PCR category") +
  theme(plot.title = element_text(hjust = 0.5, size = 35)) + scale_fill_manual(values = c("darkorange", 'chartreuse','darkslateblue', "deeppink3"))
venn4 <- ggvenn(
    four,
    set_name_size = 10, text_size = 15, fill_alpha = 0.2, stroke_size = 0.5, label_sep = "\n") + ggtitle("Shared genes 125 ng/ LOW PCR category") + 
  theme(plot.title = element_text(hjust = 0.5, size = 35)) + scale_fill_manual(values = c("darkorange", 'chartreuse','darkslateblue', "deeppink3"))

#plot_grid(venn4, venn3, venn2, venn1, ncol = 2, align="v", rel_widths = c(1, 1))

venn1
venn2
venn3
venn4
```


##Figure 11
```{r , fig.width = 10, fig.height=7, echo=FALSE}
library(cartography)
columns_of_interest = colnames(dedupCounts)[!grepl("^0_", colnames(dedupCounts))]
countsPCA = dedupCounts[, columns_of_interest]
readCds <- DGEList(counts = countsPCA, group = factor(sampleInfo$amountFac[!grepl("^NC", sampleInfo$amountFac)], levels = c("1", "2", "4", "7","15","31", "62", "125", "250", "500","1000")), samples = sampleInfo$Id[!grepl("^0_", sampleInfo$Id)])
readCds <- calcNormFactors(readCds, method = "TMM")
readSf <- 1 / (readCds$samples$norm.factors * readCds$samples$lib.size)
readSf <- readSf / ezGeomean(readSf)
readLogSig <- log2(ezScaleColumns(countsPCA, readSf) + 1)
umap = umap(t(readLogSig), n_components = 2, random_state = 15) 
layout <- umap[["layout"]] 
layout <- data.frame(layout) 
umapx = merge(layout, sampleInfo, by.x = "row.names", by.y = "Id")

colors2 <- carto.pal(pal1 = "turquoise.pal" ,n1 = 11)

ggplot(umapx, aes(X1, X2, color = amountFac, shape = Sequencer)) + 
  geom_point(aes(size = `Cycles Category`)) + 
  scale_color_manual(values = carto.pal(pal1 = "turquoise.pal" ,n1 = 11)) + 
  theme(legend.text = element_text(size = 11)) + 
  guides(shape = guide_legend(override.aes = list(size = 5)), color = guide_legend(override.aes = list(size = 5))) + 
  labs (color = "Amount", size = "PCR Cycles Category", shape = "Sequencer")
```

##Figure 12
```{r , fig.align = "center", fig.width = 16, fig.height=6, echo=FALSE, out.width='\\textwidth'}
logRatio <- sweep(readLogSig, 1, rowMeans(readLogSig))
row_sds <- apply(readLogSig, 1, sd)
top_2000_indices <- order(row_sds, decreasing = TRUE)[1:2000]
logRatio <- logRatio[top_2000_indices, ]
lim <- c(-6, 6)
design <- ezFrame(sampleInfo[ sampleInfo$amountFac != "NC", c("amountFac", "Cycles Category", "Sequencer", "Id")])
rownames(design) = design$Id
design$CyclesCategory = factor(design$'Cycles Category', levels = c("LOW", "INTERMEDIATE", "HIGH"))
design$Amount = factor(design$amount,levels= c("1", "2", "4", "7", "15", "31", "62", "125", "250", "500", "1000"))
design = design[order(design$Amount) , ]
design = design[,-1]

colors2 <- carto.pal(pal1 = "turquoise.pal" ,n1 = 11)
ann_colors <- list(Sequencer = c("Aviti" = "chartreuse", 
                                 "G4" = "darkslateblue",
                                 "Novaseq6000" = "darkorange",
                                 "NovaseqX" = "deeppink3"),
                   CyclesCategory = c("LOW" = "skyblue",
                                      "INTERMEDIATE" = "yellowgreen",
                                      "HIGH" = "orangered"), 
                   Amount = c("1" = colors2[1],
                              "2" = colors2[2],
                              "4" = colors2[3],
                              "7" = colors2[4],
                              "15" = colors2[5],
                              "31" = colors2[6],
                              "62" = colors2[7],
                              "125" = colors2[8],
                              "250" = colors2[9],
                              "500" = colors2[10],
                              "1000" = colors2[11])) 
#GCTable[ , df[with(df, order(amountFac, `Cycles Category`, Sequencer)),]$Id]
p <- pheatmap(logRatio[, df[with(df, order(amountFac, `Cycles Category`, Sequencer)),]$Id], color=getBlueRedScale(), clustering_method="ward.D2",
              breaks=seq(from=lim[1], to=lim[2], length.out=257),
              scale="row", 
              cluster_rows=TRUE,
              cluster_cols=FALSE,
              show_rownames=FALSE, 
              show_colnames=FALSE,
              annotation_col = design[,c(-1,-3)],
              annotation_colors = ann_colors
)

```

##Figure 13

### Cycles - Prededuplication
```{r , fig.width = 15, fig.height=15, echo=FALSE}
comb_allseq = ezFrame(rawCounts)
comb_allseq = comb_allseq[,colnames(comb_allseq)[!grepl("^0_", colnames(comb_allseq))]]
comb_allseq = melt(comb_allseq %>% rownames_to_column()) %>% separate(variable, into = c("amount", "cycles", "Sequencer"), sep = "_") %>% rename("Gene_names" = "rowname")

comb_allseq$cycles <- as.integer(comb_allseq$cycles)
comb_allseq$amount = factor(comb_allseq$amount, levels = c("1", "2", "4", "7", "15", "31", "62", "125", "250", "500", "1000"))

par(mfrow=c(3,4))

for (i in levels(comb_allseq$amount)) {
  
  cycles_max <- max(unique(comb_allseq[comb_allseq$amount == i,]$cycles))
  cycles_min <- min(unique(comb_allseq[comb_allseq$amount == i,]$cycles))
  X <- comb_allseq[comb_allseq$amount == i & comb_allseq$cycles %in% c(cycles_max, cycles_min),]
  X <- merge(X[X$cycles == cycles_max,] %>% dplyr::select(value, amount, Sequencer, Gene_names) %>% rename("value_HC" = "value"), X[X$cycles == cycles_min,] %>% dplyr::select(value, amount, Sequencer, Gene_names) %>% rename("value_LC" = "value"), by = c("Gene_names", "amount", "Sequencer"))
  
  plot(X$value_HC + 5, X$value_LC + 5, log = "xy", pch = 19, col = c('chartreuse','darkslateblue', "darkorange","deeppink3"), xlab = "HIGH", ylab = "LOW", main = i, cex.main = 3, cex.lab= 1.5)
  legend("topleft", legend = levels(factor(X$Sequencer)), pch = 19, col = c('chartreuse','darkslateblue', "darkorange","deeppink3"), cex = 1.5)
  x = cor.test(X$value_HC, X$value_LC,  method = "pearson")
  legend(x='bottomright', legend=paste('R=', round(x$estimate,4), ", P-value = ", round(x$p.value,4)), cex = 1.3)
}

```

### Cycles - Post deduplication
```{r , fig.width = 15, fig.height=15, echo=FALSE}
comb_allseq = ezFrame(dedupCounts)
comb_allseq = comb_allseq[,colnames(comb_allseq)[!grepl("^0_", colnames(comb_allseq))]]
comb_allseq = melt(comb_allseq %>% rownames_to_column()) %>% separate(variable, into = c("amount", "cycles", "Sequencer"), sep = "_") %>% rename("Gene_names" = "rowname")

comb_allseq$cycles <- as.integer(comb_allseq$cycles)
comb_allseq$amount = factor(comb_allseq$amount, levels = c("1", "2", "4", "7", "15", "31", "62", "125", "250", "500", "1000"))

par(mfrow=c(3,4))

for (i in levels(comb_allseq$amount)) {
  
  cycles_max <- max(unique(comb_allseq[comb_allseq$amount == i,]$cycles))
  cycles_min <- min(unique(comb_allseq[comb_allseq$amount == i,]$cycles))
  X <- comb_allseq[comb_allseq$amount == i & comb_allseq$cycles %in% c(cycles_max, cycles_min),]
  X <- merge(X[X$cycles == cycles_max,] %>% dplyr::select(value, amount, Sequencer, Gene_names) %>% rename("value_HC" = "value"), X[X$cycles == cycles_min,] %>% dplyr::select(value, amount, Sequencer, Gene_names) %>% rename("value_LC" = "value"), by = c("Gene_names", "amount", "Sequencer"))
  
  plot(X$value_HC + 5, X$value_LC + 5, log = "xy", pch = 19, col = c('chartreuse','darkslateblue', "darkorange","deeppink3"), xlab = "HIGH", ylab = "LOW", main = i, cex.main = 3, cex.lab= 1.5)
  legend("topleft", legend = levels(factor(X$Sequencer)), pch = 19, col = c('chartreuse','darkslateblue', "darkorange","deeppink3"), cex = 1.5)
  x = cor.test(X$value_HC, X$value_LC,  method = "pearson")
  legend(x='bottomright', legend=paste('R=', round(x$estimate,4), ", P-value = ", round(x$p.value,4)), cex = 1.3)
}

```

##Figure 14
```{r , fig.width = 10, fig.height=5, echo=FALSE}
percTop <- as.data.frame(apply(dedupCounts, 2, function(x) (sum(sort(x, decreasing = TRUE)[1:20]) / sum(x))* 100)) %>% rownames_to_column()
colnames(percTop) = c("Sample", "PercTopCounts")
percTop = percTop %>% separate(Sample, into = c("Amount", "cycles", "Sequencer"), sep = "_")
percTop = merge(percTop, sampleInfo, by = c("Amount", "cycles", "Sequencer"))
percTop = percTop[percTop$Amount != "0",]
percTop$Amount = factor(percTop$Amount, levels = c("1", "2", "4", "7", "15", "31", "62", "125", "250", "500", "1000"))

ggplot(percTop, aes(Amount, PercTopCounts, color = `Cycles Category`, shape = Sequencer)) + 
  geom_point(size = 2.5) + 
  scale_color_manual(values = c("skyblue", "yellowgreen", "orangered")) + 
  labs(x = "Amount", y = "Percentage of reads in top 20 genes", color = "PCR Cycles Category")

```


##Figure 15
```{r , fig.width = 15, fig.height=5, echo=FALSE}
dataInfoSorted <- dataInfo[dataInfo$Amount != "0",]
dataInfoSorted <- dataInfoSorted[ order(dataInfoSorted$Amount, dataInfoSorted$`Cycles Category`), ]
idList <- split(dataInfoSorted$Id, paste(dataInfoSorted$Amount, dataInfoSorted$`Cycles Category`))
mySet <- names(idList)[1]

corrBySample <- sapply(idList, function(idx){
  dedupCorr <- cor(dedupCounts[isAllPresent ,idx], method = "spearman")
  dedupCorr[upper.tri(dedupCorr)]
})

xLong <- reshape2::melt(corrBySample)
xLong$Var2 <- factor(xLong$Var2, levels=unique(paste(dataInfoSorted$Amount, dataInfoSorted$`Cycles Category`)))
xLong1 <- ezFrame(xLong, type="unique")

corrBySample <- sapply(idList, function(idx){
  rawCorr <- cor(rawCounts[isAllPresent,idx], method = "spearman")
  rawCorr[upper.tri(rawCorr)]
})

xLong <- reshape2::melt(corrBySample)
xLong$Var2 <- factor(xLong$Var2, levels=unique(paste(dataInfoSorted$Amount, dataInfoSorted$`Cycles Category`)))
xLong2 <- ezFrame(xLong, type="raw")
xLong <- rbind(xLong1, xLong2)

xLong <- xLong %>% separate(Var2, c("amountFac", "Cycles Category"), " ")
xLong$amountFac <- factor(xLong$amountFac, levels=c("1", "2", "4", "7", "15", "31", "62", "125", "250", "500", "1000"))
xLong$`Cycles Category` = factor(xLong$`Cycles Category`, levels = c("LOW", "INTERMEDIATE", "HIGH"))

ggplot(xLong %>% group_by(amountFac, `Cycles Category`, type) %>% summarise(median = median(value)), 
       aes(x = amountFac, y = median, col = type, group = type)) + 
    labs(x = "Amount", y = "Median Correlation", color = "Status") +
    facet_wrap(~`Cycles Category`) + 
    geom_point(size = 3) + geom_line() + 
    theme(axis.text.x = element_text(size = 10), text = element_text(size = 17)) +
    scale_color_manual(values=c('Purple','Turquoise'), labels=c('Raw', 'Unique')) + 
    guides( color = guide_legend(override.aes = list(size = 5))) 
```


##Figure 16
```{r , fig.width = 10, fig.height=5, echo=FALSE}
#Run qualimap
# resBaseDir <- "/srv/GT/analysis/p2220/PCR-duplicates-study/qualimap-dedup"
# if (!file.exists(resBaseDir)){
#   dir.create(resBaseDir, recursive = TRUE)
# }
# 
 # for (seqr in names(dedupFolders)){
 #   bamFiles <- list.files(dedupFolders[seqr], pattern=".*dedup.bam", full.names=TRUE)
 #   sampleNames <- sub(".dedup.bam", "", basename(bamFiles))
 #   sampleNames <- sub("_S.*", "", sampleNames)
 #   names(bamFiles) <- sampleNames
 #   for (sm in unique(sampleInfo$Sample)){
 #     resDir <- paste0(resBaseDir, "/", sm, "_", seqr)
 #     message(resDir)
 #     cmd <- paste("unset DISPLAY; /usr/local/ngseq/packages/QC/Qualimap/2.2.1/qualimap bamqc -bam", bamFiles[sm], "--java-mem-size=4G", "-c -nw 400 -hm 3 -nt 8", "-outdir", resDir)
 #     ezSystem(cmd)
 #   }
 # }


files = list.files("/srv/GT/analysis/p2220/PCR-duplicates-study/qualimap_NovaseqX100bp_dedup/", pattern = "mapped_reads_gc-content_distribution.txt", recursive = TRUE)
files = paste0("/srv/GT/analysis/p2220/PCR-duplicates-study/qualimap_NovaseqX100bp_dedup/", files)
sample_names = list.files("/srv/GT/analysis/p2220/PCR-duplicates-study/qualimap_NovaseqX100bp_dedup/")
GCfiles = data.frame("File" = files, "Sample" = sample_names)

GCTable = read_delim(GCfiles$File[1])
colnames(GCTable) = c("GC_Content", GCfiles[GCfiles$File == GCfiles$File[1],]$Sample)
for (i in GCfiles$File[-1]){
  name = GCfiles[GCfiles$File == i,]$Sample
  x = read_delim(i)
  colnames(x) = c("GC_Content", name)
  GCTable = merge(GCTable, x, by = "GC_Content")
}

GCTableM = melt(GCTable, id.vars = "GC_Content") %>% separate(variable, into = c("Amount", "cycles", "Sequencer"))
GCTableM = merge(GCTableM, sampleInfo, by = c("Amount", "cycles", "Sequencer"))
GCTableM = GCTableM[GCTableM$Amount != "0",]
GCTableM$amountFac = factor(GCTableM$amountFac, levels = levels(GCTableM$amountFac)[-1])

ggplot(GCTableM, aes(GC_Content, value, color = Sequencer)) + geom_line() + facet_grid(amountFac ~`Cycles Category`) + labs(x = "GC content %", y = "Fraction of reads", color = "Amount")

```


```{r , fig.width = 11, fig.height=6, echo=FALSE}
design = ezFrame(dataInfo[ , c("amountFac", "Cycles Category", "Sequencer")], row.names = dataInfo$Id)
colnames(design) = c("Amount", "CyclesCategory", "Sequencer")

GCTable = GCTable[, colnames(GCTable)[!grepl("^0_", colnames(GCTable))]]
design = design[design$Amount != "NC",]
design$Amount = factor(design$Amount, levels = levels(design$Amount)[-1])

colors2 <- carto.pal(pal1 = "turquoise.pal" , n1 = 11)
ann_colors <- list(Sequencer = c("Aviti" = "chartreuse", 
                                 "G4" = "darkslateblue",
                                 "Novaseq6000" = "darkorange",
                                 "NovaseqX" = "deeppink3"),
                   CyclesCategory = c("LOW" = "skyblue",
                                      "INTERMEDIATE" = "yellowgreen",
                                      "HIGH" = "orangered"), 
                   Amount = c("1" = colors2[1],
                              "2" = colors2[2],
                              "4" = colors2[3],
                              "7" = colors2[4],
                              "15" = colors2[5],
                              "31" = colors2[6],
                              "62" = colors2[7],
                              "125" = colors2[8],
                              "250" = colors2[9],
                              "500" = colors2[10],
                              "1000" = colors2[11])) 
df = dataInfo[dataInfo$amountFac != "NC",]
pheatmap(log2(GCTable[ , df[with(df, order(amountFac, `Cycles Category`, Sequencer)),]$Id] + 0.02), 
         annotation_col=design, 
         scale="none", 
         cluster_rows=FALSE,
         cluster_cols=FALSE,
         show_rownames=TRUE, show_colnames = FALSE,
         annotation_colors = ann_colors, fontsize = 6
         )
setHook("grid.newpage", NULL, "replace")
grid.text("GC content", x = 0.85 , rot=-90, gp=gpar(fontsize=16))
```
````
