## Supplementary material for "The impact of PCR duplication on RNAseq data generated using NovaSeq 6000, NovaSeq X, AVITI and G4 sequencers": braken.sh

```
for i in $(ls *.report.txt);
do
	python /usr/local/ngseq/src/Bracken/src/est_abundance.py -i ${i} -k /srv/GT/databases/kraken2/standard_20230605/database100mers.kmer_distrib -l S -t 10 -o ${i}.bracken;
done
for i in $(ls *.fastq.gz.txt); do awk '$3 == 0 {print}' $i | wc -l >> unclassified.txt; done
for i in $(ls *.fastq.gz.txt); do awk '$3 != 0 {print}' $i | wc -l >> classified.txt; done
for i in $(ls *.fastq.gz.txt); do echo $i >> sampleid.txt; done
paste -d"\t" sampleid.txt classified.txt unclassified.txt | sed 's/.markedUMI.subsampled.fastq.gz.txt//g' | sed 1i'sampleid classified unclassified' > kraken_read_summary.txt
```
