## Supplementary material for "The impact of PCR duplication on RNAseq data generated using NovaSeq 6000, NovaSeq X, AVITI and G4 sequencers": featurecounts.R

```
library(Rsubread)

filenames <- list.files(path = "/srv/GT/analysis/zajacn/p2220/o31444_Aviti/Aviti_p2220_o31444_subsampled_dedup/", pattern = ".dedup.bam")
gtffile <- file.path("/srv/GT/reference/Homo_sapiens/GENCODE/GRCh38.p13/Annotation/Release_42-2023-01-30/Genes/genes.gtf")
post <- featureCounts(files=paste0("/srv/GT/analysis/zajacn/p2220/o31444_Aviti/Aviti_p2220_o31444_subsampled_dedup/",filenames), 
                    annot.ext=gtffile, 
                    isGTFAnnotationFile=TRUE,
                    isPairedEnd=FALSE, 
                    allowMultiOverlap=TRUE, 
                    strandSpecific=2, 
                    minOverlap=10, 
                    ignoreDup=TRUE, 
                    countMultiMappingReads=TRUE, 
                    fraction=TRUE, 
                    primaryOnly=FALSE)
colnames(post$counts) <- str_replace(colnames(post$counts), "_S[0-9]+_L001.dedup.bam", ".dedup.bam")
saveRDS(post, "/srv/GT/analysis/zajacn/p2220/o31444_Aviti/Aviti_p2220_o31444_subsampled_dedup/featurecounts.table.POST.rds")

filenames <- list.files(path = "/srv/gstore/projects/p2220/STAR_markdups_Aviti_subsampled_2023-08-24--10-09-41/", pattern = ".bam$")
pre <- featureCounts(files=paste0("/srv/gstore/projects/p2220/STAR_markdups_Aviti_subsampled_2023-08-24--10-09-41/",filenames), 
                    annot.ext=gtffile, 
                    isGTFAnnotationFile=TRUE,
                    isPairedEnd=FALSE, 
                    allowMultiOverlap=TRUE, 
                    strandSpecific=2, 
                    minOverlap=10, 
                    ignoreDup=TRUE, 
                    countMultiMappingReads=TRUE, 
                    fraction=TRUE, 
                    primaryOnly=FALSE)
colnames(pre$counts) <- str_replace(colnames(pre$counts), "_S[0-9]+_L001.bam", ".bam")
saveRDS(pre, "/srv/GT/analysis/zajacn/p2220/o31444_Aviti/Aviti_p2220_o31444_subsampled_dedup/featurecounts.table.PRE.rds")
pre_post <- merge(pre$counts, post$counts, by = 'row.names', all = TRUE)
rownames(pre_post) <- pre_post$Row.names
pre_post <- pre_post[,-1]
colnames(pre_post) <- str_replace(str_replace(colnames(pre_post), ".bam", "_pre"), ".dedup_pre", "_post")
write.table(pre_post, "/srv/GT/analysis/zajacn/p2220/o31444_Aviti/Aviti_p2220_o31444_subsampled_dedup/featurecounts.pre.post.combined.txt", sep = "\t")
```
