## Supplementary material for "The impact of PCR duplication on RNAseq data generated using NovaSeq 6000, NovaSeq X, AVITI and G4 sequencers": kraken.sh

```
#!/bin/bash
#SBATCH --partition=employee
#SBATCH --array=1-36
#SBATCH --cpus-per-task=1
#SBATCH --time=1-00:00:00
#SBATCH --error=/srv/GT/analysis/zajacn/p2220/o31444_Aviti/Aviti_p2220_o31444_subsampled_dedup/contamination/cont.%J.err
#SBATCH --output=/srv/GT/analysis/zajacn/p2220/o31444_Aviti/Aviti_p2220_o31444_subsampled_dedup/contamination/cont.%J.out
#SBATCH --mem-per-cpu=160G
#SBATCH --job-name=p2220

source /usr/local/ngseq/etc/lmod_profile
module load Tools/kraken/2.0.9

DIR=/srv/GT/analysis/zajacn/p2220/o31444_Aviti/Aviti_p2220_o31444_subsampled_dedup/contamination

k=$SLURM_ARRAY_TASK_ID
name=`sed -n ${k}p < /srv/GT/analysis/zajacn/p2220/o31444_Aviti/Aviti_p2220_o31444_subsampled_dedup/contamination/list_of_files`

kraken2 -db /srv/GT/databases/kraken2/standard_20230605/ --confidence 0 --minimum-base-quality 0 --output $DIR/${name}.txt --report $DIR/${name}.report.txt --threads 8 /srv/GT/analysis/zajacn/p2220/o31444_Aviti/Aviti_p2220_o31444_subsampled/FastP/${name}  1>  $DIR/${name}.log
```
