## Supplementary material for "The impact of PCR duplication on RNAseq data generated using NovaSeq 6000, NovaSeq X, AVITI and G4 sequencers": seqtk.sh

```
#!/bin/bash
#SBATCH --partition=employee
#SBATCH --array=1-36
#SBATCH --cpus-per-task=1
#SBATCH --time=1-00:00:00
#SBATCH --error=/srv/GT/analysis/zajacn/p2220/o31444_Aviti/Aviti_p2220_o31444_subsampled/subsample.%J.err
#SBATCH --output=/srv/GT/analysis/zajacn/p2220/o31444_Aviti/Aviti_p2220_o31444_subsampled/subsample.%J.out
#SBATCH --mem-per-cpu=20G
#SBATCH --job-name=p2220

source /usr/local/ngseq/etc/lmod_profile
module load Tools/seqtk/1.3

DIR=/srv/GT/analysis/zajacn/p2220/o31444_Aviti/Aviti_p2220_o31444_subsampled

k=$SLURM_ARRAY_TASK_ID
name=`sed -n ${k}p < /srv/GT/analysis/zajacn/p2220/o31444_Aviti/list_of_files`

seqtk sample -s100 /srv/gstore/projects/p2220/Aviti_p2220_o31444/${name} 2000000 > $DIR/${name}.subsampled.fastq
gzip $DIR/${name}.subsampled.fastq
```
