## Supplementary material for "The impact of PCR duplication on RNAseq data generated using NovaSeq 6000, NovaSeq X, AVITI and G4 sequencers": Map_0_17_S1_L001_Aviti_p2220_o31444_subsampled.sh

```
#!/bin/bash

set -e
set -o pipefail
umask 0002

#### SET THE STAGE
SCRATCH_DIR=/scratch/STAR_markdups_Aviti_subsampled_2023-08-24--10-09-41_0_17_S1_L001_temp$$
GSTORE_DIR=/srv/gstore/projects
INPUT_DATASET=/srv/gstore/projects/p2220/STAR_markdups_Aviti_subsampled_2023-08-24--10-09-41/input_dataset.tsv
LAST_JOB=FALSE
echo "Job runs on `hostname`"
echo "at $SCRATCH_DIR"
mkdir $SCRATCH_DIR || exit 1
cd $SCRATCH_DIR || exit 1
source /usr/local/ngseq/etc/lmod_profile
module add Aligner/STAR/2.7.10b Tools/samtools/1.17 Dev/jdk/14 Dev/R/4.3.0 Dev/Python/3.10.9 Tools/Picard/2.22.8 QC/fastp/0.23.4

#### NOW THE ACTUAL JOBS STARTS
R --vanilla --slave<<  EOT
EZ_GLOBAL_VARIABLES <<- '/usr/local/ngseq/opt/EZ_GLOBAL_VARIABLES.txt'
library(ezRun)
param = list()
param[['cores']] = '8'
param[['ram']] = '30'
param[['scratch']] = '100'
param[['partition']] = 'employee'
param[['process_mode']] = 'SAMPLE'
param[['samples']] = '0_17_S1_L001,0_19_S2_L001,0_21_S3_L001,1_16_S13_L001,1_18_S14_L001,1_20_S15_L001,1000_10_S4_L001,1000_6_S5_L001,1000_8_S6_L001,125_11_S7_L001,125_13_S8_L001,125_9_S9_L001,15_12_S10_L001,15_14_S11_L001,15_16_S12_L001,2_15_S19_L001,2_17_S20_L001,2_19_S21_L001,250_10_S16_L001,250_12_S17_L001,250_8_S18_L001,31_11_S22_L001,31_13_S23_L001,31_15_S24_L001,4_14_S25_L001,4_16_S26_L001,4_18_S27_L001,500_11_S28_L001,500_7_S29_L001,500_9_S30_L001,62_10_S31_L001,62_12_S32_L001,62_14_S33_L001,7_13_S34_L001,7_15_S35_L001,7_17_S36_L001'
param[['refBuild']] = 'Homo_sapiens/GENCODE/GRCh38.p13/Annotation/Release_42-2023-01-30'
param[['paired']] = 'false'
param[['strandMode']] = 'antisense'
param[['refFeatureFile']] = 'genes.gtf'
param[['controlSeqs']] = ''
param[['cmdOptions']] = '--sjdbOverhang 150 --outFilterType BySJout --outFilterMatchNmin 30 --outFilterMismatchNmax 10 --outFilterMismatchNoverLmax 0.05 --outMultimapperOrder Random --alignSJDBoverhangMin 1 --alignSJoverhangMin 8 --alignIntronMax 100000 --alignMatesGapMax 100000  --outFilterMultimapNmax 50 --chimSegmentMin 15 --chimJunctionOverhangMin 15 --chimScoreMin 15 --chimScoreSeparation 10 --outSAMstrandField intronMotif --alignEndsProtrude 3 ConcordantPair'
param[['getJunctions']] = 'false'
param[['twopassMode']] = 'false'
param[['trimAdapter']] = 'true'
param[['trim_front1']] = '0'
param[['trim_tail1']] = '0'
param[['cut_front']] = 'false'
param[['cut_front_window_size']] = '4'
param[['cut_front_mean_quality']] = '20'
param[['cut_tail']] = 'false'
param[['cut_tail_window_size']] = '4'
param[['cut_tail_mean_quality']] = '20'
param[['cut_right']] = 'false'
param[['cut_right_window_size']] = '4'
param[['cut_right_mean_quality']] = '20'
param[['average_qual']] = '0'
param[['max_len1']] = '0'
param[['max_len2']] = '0'
param[['poly_x_min_len']] = '10'
param[['length_required']] = '18'
param[['cmdOptionsFastp']] = ''
param[['markDuplicates']] = 'false'
param[['specialOptions']] = ''
param[['mail']] = ''
param[['sushi_app']] = 'STARApp'
param[['nice']] = '100'
param[['dataRoot']] = '/srv/gstore/projects'
param[['resultDir']] = 'p2220/STAR_markdups_Aviti_subsampled_2023-08-24--10-09-41'
param[['isLastJob']] = FALSE
output = list()
output[['Name']] = '0_17_S1_L001'
output[['BAM [File]']] = 'p2220/STAR_markdups_Aviti_subsampled_2023-08-24--10-09-41/0_17_S1_L001.bam'
output[['BAI [File]']] = 'p2220/STAR_markdups_Aviti_subsampled_2023-08-24--10-09-41/0_17_S1_L001.bam.bai'
output[['IGV [Link]']] = 'p2220/STAR_markdups_Aviti_subsampled_2023-08-24--10-09-41/0_17_S1_L001-igv.html'
output[['Species']] = ' '
output[['refBuild']] = 'Homo_sapiens/GENCODE/GRCh38.p13/Annotation/Release_42-2023-01-30'
output[['paired']] = 'false'
output[['refFeatureFile']] = 'genes.gtf'
output[['strandMode']] = 'antisense'
output[['Read Count']] = ''
output[['IGV [File]']] = 'p2220/STAR_markdups_Aviti_subsampled_2023-08-24--10-09-41/0_17_S1_L001-igv.html'
output[['PreprocessingLog [File]']] = 'p2220/STAR_markdups_Aviti_subsampled_2023-08-24--10-09-41/0_17_S1_L001_preprocessing.log'
output[['STARLog [File]']] = 'p2220/STAR_markdups_Aviti_subsampled_2023-08-24--10-09-41/0_17_S1_L001_STAR.log'
input = list()
input[['Name']] = '0_17_S1_L001'
input[['Read1']] = 'p2220/Aviti_p2220_o31444_subsampled/0_17_S1_L001_R1_001.subsampled.fastq.gz'
input[['Read Number']] = '1128694'
input[['Species']] = ' '
EzAppSTAR\$new()\$run(input=input, output=output, param=param)
EOT


#### JOB IS DONE WE PUT THINGS IN PLACE AND CLEAN AUP
. "/usr/local/ngseq/miniconda3/etc/profile.d/conda.sh"
conda activate sushi
g-req -w copy 0_17_S1_L001.bam 0_17_S1_L001.bam.bai 0_17_S1_L001-igv.html 0_17_S1_L001_preprocessing.log 0_17_S1_L001_STAR.log /srv/gstore/projects/p2220/STAR_markdups_Aviti_subsampled_2023-08-24--10-09-41
cd /scratch
rm -rf /scratch/STAR_markdups_Aviti_subsampled_2023-08-24--10-09-41_0_17_S1_L001_temp$$ || exit 1
```
