## Supplementary material for "The impact of PCR duplication on RNAseq data generated using NovaSeq 6000, NovaSeq X, AVITI and G4 sequencers": dedup.sh

```
#!/bin/bash
#SBATCH --partition=employee
#SBATCH --array=1-36
#SBATCH --cpus-per-task=20
#SBATCH --time=1-00:00:00
#SBATCH --mem-per-cpu=15G
#SBATCH --job-name=p2220_dedup

source /usr/local/ngseq/miniconda3/etc/profile.d/conda.sh
conda activate umi_tools

DIR=/srv/GT/analysis/zajacn/p2220/o31444_G4/G4_p2220_o31444_subsampledv2_dedup
k=$SLURM_ARRAY_TASK_ID
name=`sed -n ${k}p < /srv/GT/analysis/zajacn/p2220/o31444_G4/G4_p2220_o31444_subsampledv2_dedup/list_of_files`

umi_tools dedup --stdin=/srv/gstore/projects/p2220/STAR_markdups_G4_subsampled_2023-08-24--10-16-09/${name}.bam --output-stats=${name} --stdout=$DIR/${name}.dedup.bam  --umi-separator=":" --log=$DIR/${name}.log
```
