## Supplementary material for "The impact of PCR duplication on RNAseq data generated using NovaSeq 6000, NovaSeq X, AVITI and G4 sequencers": braken.sh

```
#!/bin/bash
#SBATCH --partition=employee
#SBATCH --cpus-per-task=1
#SBATCH --time=1-00:00:00
#SBATCH --error=/srv/GT/analysis/zajacn/p2220/o31444/o31444_NovaSeq_NOV1703_subsampled_dedup/contamination/brak.err
#SBATCH --output=/srv/GT/analysis/zajacn/p2220/o31444/o31444_NovaSeq_NOV1703_subsampled_dedup/contamination/brak.out
#SBATCH --mem-per-cpu=250G
#SBATCH --job-name=braken

source /usr/local/ngseq/etc/lmod_profile
module load Tools/kraken/2.0.9
module load Dev/Python/3.9.7

for i in $(ls *.report.txt);
do
	python /usr/local/ngseq/src/Bracken/src/est_abundance.py -i ${i} -k /srv/GT/databases/kraken2/standard_20230605/database100mers.kmer_distrib -l S -t 10 -o ${i}.bracken; #changed to -l G
done
for i in $(ls *.fastq.gz.txt); do awk '$3 == 0 {print}' $i | wc -l >> unclassified.txt; done
for i in $(ls *.fastq.gz.txt); do awk '$3 != 0 {print}' $i | wc -l >> classified.txt; done
for i in $(ls *.fastq.gz.txt); do echo $i >> sampleid.txt; done
paste -d"\t" sampleid.txt classified.txt unclassified.txt | sed 's/.markedUMI.subsampled.fastq.gz.txt//g' | sed 1i'sampleid classified unclassified' > kraken_read_summary.txt
```
