## Supplementary material for "The impact of PCR duplication on RNAseq data generated using NovaSeq 6000, NovaSeq X, AVITI and G4 sequencers": dedup.sh

```
#!/bin/bash
#SBATCH --partition=employee
#SBATCH --array=1-36
#SBATCH --cpus-per-task=20
#SBATCH --time=1-00:00:00
#SBATCH --mem-per-cpu=15G
#SBATCH --job-name=p2220_dedup

source /usr/local/ngseq/miniconda3/etc/profile.d/conda.sh
conda activate umi_tools

DIR=/srv/GT/analysis/zajacn/p2220/o31444/o31444_NovaSeq_NOV1703_subsampled_dedup
k=$SLURM_ARRAY_TASK_ID
name=`sed -n ${k}p < /srv/GT/analysis/zajacn/p2220/o31444/o31444_NovaSeq_NOV1703_dedup/list_of_files_interest`

umi_tools dedup --stdin=/srv/gstore/projects/p2220/o31444_STAR_markdups_Novaseq_subsampled_2023-08-24--10-13-19/${name}.bam --output-stats=${name} --stdout=$DIR/${name}.dedup.bam  --log=${name}.log
```
