## Supplementary material for "The impact of PCR duplication on RNAseq data generated using NovaSeq 6000, NovaSeq X, AVITI and G4 sequencers": extractUMI.sh

```
for i in $(ls ../*.fastq.gz| sed 's/_R[1-2]_001.fastq.gz//g' | sed 's"../""g' | sort | uniq); do umi_tools extract --stdin=../${i}_R2_001.fastq.gz --read2-in=../${i}_R1_001.fastq.gz --stdout=discard.fastq --read2-out=${i}.markedUMI.fastq.gz --bc-pattern=NNNNNNNNNNN ; done
```
