## Supplementary material for "The impact of PCR duplication on RNAseq data generated using NovaSeq 6000, NovaSeq X, AVITI and G4 sequencers": fastp.sh

```
#!/bin/bash
#SBATCH --partition=employee
#SBATCH --array=1-36
#SBATCH --cpus-per-task=20
#SBATCH --time=1-00:00:00
#SBATCH --mem-per-cpu=15G
#SBATCH --job-name=p2220_fastp

module load QC/fastp/0.23.4

DIR2=/srv/GT/analysis/zajacn/p2220/o31444_NovaseqX_100bp/FastP
DIR=/srv/GT/analysis/zajacn/p2220/o31444_NovaseqX/o31444_NovaSeq_230825_X003_markedUMIs_subsampled
k=$SLURM_ARRAY_TASK_ID
name=`sed -n ${k}p < /srv/GT/analysis/zajacn/p2220/o31444_NovaseqX/o31444_NovaSeq_230825_X003_markedUMIs_subsampled/list_of_files`
ADAPTERS=/srv/GT/databases/contaminants/allIllumina-forTrimmomatic-20160202.fa

fastp --in1 $DIR/${name} --out1 $DIR2/${name} --thread 8 --trim_front1 0 --trim_tail1 50 --average_qual 0 --adapter_fasta $ADAPTERS --max_len1 0 --max_len2 0 --trim_poly_x --poly_x_min_len 10 --length_required 18 --compression 4 2> ${name}_preprocessing.log
```
