## Supplementary material for "The impact of PCR duplication on RNAseq data generated using NovaSeq 6000, NovaSeq X, AVITI and G4 sequencers": Map_0_17_o31444_NovaSeq_230825_X003_markedUMIs_subsampled.sh

```
#!/bin/bash

set -e
set -o pipefail
umask 0002

#### SET THE STAGE
SCRATCH_DIR=/scratch/o31444_STAR_cutto100bp_2023-12-07--17-39-06_0_17_temp$$
GSTORE_DIR=/srv/gstore/projects
INPUT_DATASET=/srv/gstore/projects/p2220/o31444_STAR_cutto100bp_2023-12-07--17-39-06/input_dataset.tsv
LAST_JOB=FALSE
echo "Job runs on `hostname`"
echo "at $SCRATCH_DIR"
mkdir $SCRATCH_DIR || exit 1
cd $SCRATCH_DIR || exit 1
source /usr/local/ngseq/etc/lmod_profile
module add Aligner/STAR/2.7.10b Tools/samtools/1.17 Dev/jdk/14 Dev/R/4.3.2 Dev/Python/3.10.9 Tools/Picard/2.22.8 QC/fastp/0.23.4

#### NOW THE ACTUAL JOBS STARTS
R --vanilla --slave<<  EOT
EZ_GLOBAL_VARIABLES <<- '/usr/local/ngseq/opt/EZ_GLOBAL_VARIABLES.txt'
if (!library(ezRun, logical.return = TRUE)){
message('retry loading ezRun')
Sys.sleep(120)
library(ezRun)
}
param = list()
param[['cores']] = '8'
param[['ram']] = '30'
param[['scratch']] = '100'
param[['partition']] = 'employee'
param[['process_mode']] = 'SAMPLE'
param[['samples']] = '0_17,0_19,0_21,1000_10,1000_6,1000_8,125_11,125_13,125_9,15_12,15_14,15_16,1_16,1_18,1_20,250_10,250_12,250_8,2_15,2_17,2_19,31_11,31_13,31_15,4_14,4_16,4_18,500_11,500_7,500_9,62_10,62_12,62_14,7_13,7_15,7_17'
param[['refBuild']] = 'Homo_sapiens/GENCODE/GRCh38.p13/Annotation/Release_42-2023-01-30'
param[['paired']] = 'false'
param[['strandMode']] = 'antisense'
param[['refFeatureFile']] = 'genes.gtf'
param[['controlSeqs']] = ''
param[['cmdOptions']] = '--sjdbOverhang 150 --outFilterType BySJout --outFilterMatchNmin 30 --outFilterMismatchNmax 10 --outFilterMismatchNoverLmax 0.05 --outMultimapperOrder Random --alignSJDBoverhangMin 1 --alignSJoverhangMin 8 --alignIntronMax 100000 --alignMatesGapMax 100000  --outFilterMultimapNmax 50 --chimSegmentMin 15 --chimJunctionOverhangMin 15 --chimScoreMin 15 --chimScoreSeparation 10 --outSAMstrandField intronMotif --alignEndsProtrude 3 ConcordantPair'
param[['getJunctions']] = 'false'
param[['twopassMode']] = 'false'
param[['trimAdapter']] = 'true'
param[['trim_front1']] = '0'
param[['trim_tail1']] = '50'
param[['cut_front']] = 'false'
param[['cut_front_window_size']] = '4'
param[['cut_front_mean_quality']] = '20'
param[['cut_tail']] = 'false'
param[['cut_tail_window_size']] = '4'
param[['cut_tail_mean_quality']] = '20'
param[['cut_right']] = 'false'
param[['cut_right_window_size']] = '4'
param[['cut_right_mean_quality']] = '20'
param[['average_qual']] = '0'
param[['max_len1']] = '0'
param[['max_len2']] = '0'
param[['poly_x_min_len']] = '10'
param[['length_required']] = '18'
param[['cmdOptionsFastp']] = ''
param[['markDuplicates']] = 'false'
param[['specialOptions']] = ''
param[['mail']] = ''
param[['sushi_app']] = 'STARApp'
param[['nice']] = '100'
param[['dataRoot']] = '/srv/gstore/projects'
param[['resultDir']] = 'p2220/o31444_STAR_cutto100bp_2023-12-07--17-39-06'
param[['isLastJob']] = FALSE
output = list()
output[['Name']] = '0_17'
output[['BAM [File]']] = 'p2220/o31444_STAR_cutto100bp_2023-12-07--17-39-06/0_17.bam'
output[['BAI [File]']] = 'p2220/o31444_STAR_cutto100bp_2023-12-07--17-39-06/0_17.bam.bai'
output[['IGV [Link]']] = 'p2220/o31444_STAR_cutto100bp_2023-12-07--17-39-06/0_17-igv.html'
output[['Species']] = 'Homo sapiens (human)'
output[['refBuild']] = 'Homo_sapiens/GENCODE/GRCh38.p13/Annotation/Release_42-2023-01-30'
output[['paired']] = 'false'
output[['refFeatureFile']] = 'genes.gtf'
output[['strandMode']] = 'antisense'
output[['Read Count']] = '2000000'
output[['IGV [File]']] = 'p2220/o31444_STAR_cutto100bp_2023-12-07--17-39-06/0_17-igv.html'
output[['PreprocessingLog [File]']] = 'p2220/o31444_STAR_cutto100bp_2023-12-07--17-39-06/0_17_preprocessing.log'
output[['STARLog [File]']] = 'p2220/o31444_STAR_cutto100bp_2023-12-07--17-39-06/0_17_STAR.log'
output[['Condition [Factor]']] = 'NA'
output[['Sample Id [B-Fabric]']] = '554700'
output[['Order Id [B-Fabric]']] = '31444'
output[['Tube [Characteristic]']] = '31444/36'
output[['PlatePosition [Characteristic]']] = 'C12'
output[['PlateName [Characteristic]']] = '2258'
input = list()
input[['Name']] = '0_17'
input[['Condition']] = 'NA'
input[['Read1']] = 'p2220/o31444_NovaSeq_230825_X003_markedUMIs_subsampled/0_17_S58.markedUMI.subsampled.fastq.gz'
input[['Species']] = 'Homo sapiens (human)'
input[['Tube']] = '31444/36'
input[['Sample Id']] = '554700'
input[['barcode1']] = 'GGAATGTC'
input[['barcode2']] = 'GTGTTCCT'
input[['LibraryPrepKit']] = 'Illumina Truseq mRNA'
input[['strandMode']] = 'antisense'
input[['Order Id']] = '31444'
input[['PlatePosition']] = 'C12'
input[['PlateName']] = '2258'
input[['Read Count']] = '2000000'
EzAppSTAR\$new()\$run(input=input, output=output, param=param)
EOT


#### JOB IS DONE WE PUT THINGS IN PLACE AND CLEAN AUP
. "/usr/local/ngseq/miniconda3/etc/profile.d/conda.sh"
conda activate sushi
g-req -w copy 0_17.bam 0_17.bam.bai 0_17-igv.html 0_17_preprocessing.log 0_17_STAR.log /srv/gstore/projects/p2220/o31444_STAR_cutto100bp_2023-12-07--17-39-06
cd /scratch
rm -rf /scratch/o31444_STAR_cutto100bp_2023-12-07--17-39-06_0_17_temp$$ || exit 1

WORKUNIT_ID=297435
update_resource_size -w $WORKUNIT_ID
```
