## Supplementary material for "The impact of PCR duplication on RNAseq data generated using NovaSeq 6000, NovaSeq X, AVITI and G4 sequencers": QC_o31444_PostDeduplication_allBams.sh

```
#!/bin/bash

set -e
set -o pipefail
umask 0002

#### SET THE STAGE
SCRATCH_DIR=/scratch/RNABamStats_2023-12-09--17-33-58_temp$$
GSTORE_DIR=/srv/gstore/projects
INPUT_DATASET=/srv/gstore/projects/p2220/RNABamStats_2023-12-09--17-33-58/input_dataset.tsv
LAST_JOB=TRUE
echo "Job runs on `hostname`"
echo "at $SCRATCH_DIR"
mkdir $SCRATCH_DIR || exit 1
cd $SCRATCH_DIR || exit 1
source /usr/local/ngseq/etc/lmod_profile
module add Tools/samtools/1.17 Dev/Python/3.10.9 Tools/Picard/2.22.8 Tools/BamUtil/1.0.14 Dev/R/4.3.2

#### NOW THE ACTUAL JOBS STARTS
R --vanilla --slave<<  EOT
EZ_GLOBAL_VARIABLES <<- '/usr/local/ngseq/opt/EZ_GLOBAL_VARIABLES.txt'
if (!library(ezRun, logical.return = TRUE)){
message('retry loading ezRun')
Sys.sleep(120)
library(ezRun)
}
param = list()
param[['cores']] = '8'
param[['ram']] = '100'
param[['scratch']] = '100'
param[['partition']] = 'employee'
param[['process_mode']] = 'DATASET'
param[['samples']] = '0_17.Aviti,0_17.G4,0_17.Novaseq6000,0_17.NovaseqX,0_19.Aviti,0_19.G4,0_19.Novaseq6000,0_19.NovaseqX,0_21.Aviti,0_21.G4,0_21.Novaseq6000,0_21.NovaseqX,1000_10.Aviti,1000_10.G4,1000_10.Novaseq6000,1000_10.NovaseqX,1000_6.Aviti,1000_6.G4,1000_6.Novaseq6000,1000_6.NovaseqX,1000_8.Aviti,1000_8.G4,1000_8.Novaseq6000,1000_8.NovaseqX'
param[['paired']] = 'false'
param[['name']] = 'RNA_BAM_Statistics'
param[['refBuild']] = 'Homo_sapiens/GENCODE/GRCh38.p13/Annotation/Release_42-2023-01-30'
param[['refFeatureFile']] = 'genes.gtf'
param[['strandMode']] = 'both'
param[['specialOptions']] = ''
param[['mail']] = ''
param[['sushi_app']] = 'RnaBamStatsApp'
param[['dataRoot']] = '/srv/gstore/projects'
param[['resultDir']] = 'p2220/RNABamStats_2023-12-09--17-33-58'
param[['isLastJob']] = TRUE
output = list()
output[['Name']] = 'RNA_BAM_Statistics'
output[['Report [File]']] = 'p2220/RNABamStats_2023-12-09--17-33-58/RNA_BAM_Statistics'
output[['Html [Link]']] = 'p2220/RNABamStats_2023-12-09--17-33-58/RNA_BAM_Statistics/00index.html'
output[['Species']] = 'Homo sapiens (human)'
output[['refBuild']] = 'Homo_sapiens/GENCODE/GRCh38.p13/Annotation/Release_42-2023-01-30'
output[['refFeatureFile']] = 'genes.gtf'
input = '/srv/gstore/projects/p2220/RNABamStats_2023-12-09--17-33-58/input_dataset.tsv'
EzAppRnaBamStats\$new()\$run(input=input, output=output, param=param)
EOT


#### JOB IS DONE WE PUT THINGS IN PLACE AND CLEAN AUP
. "/usr/local/ngseq/miniconda3/etc/profile.d/conda.sh"
conda activate sushi
g-req -w copy RNA_BAM_Statistics /srv/gstore/projects/p2220/RNABamStats_2023-12-09--17-33-58
cd /scratch
rm -rf /scratch/RNABamStats_2023-12-09--17-33-58_temp$$ || exit 1
```
