## Supplementary Figure 1 for "The impact of PCR duplication on RNAseq data generated using NovaSeq 6000, NovaSeq X, AVITI and G4 sequencers"

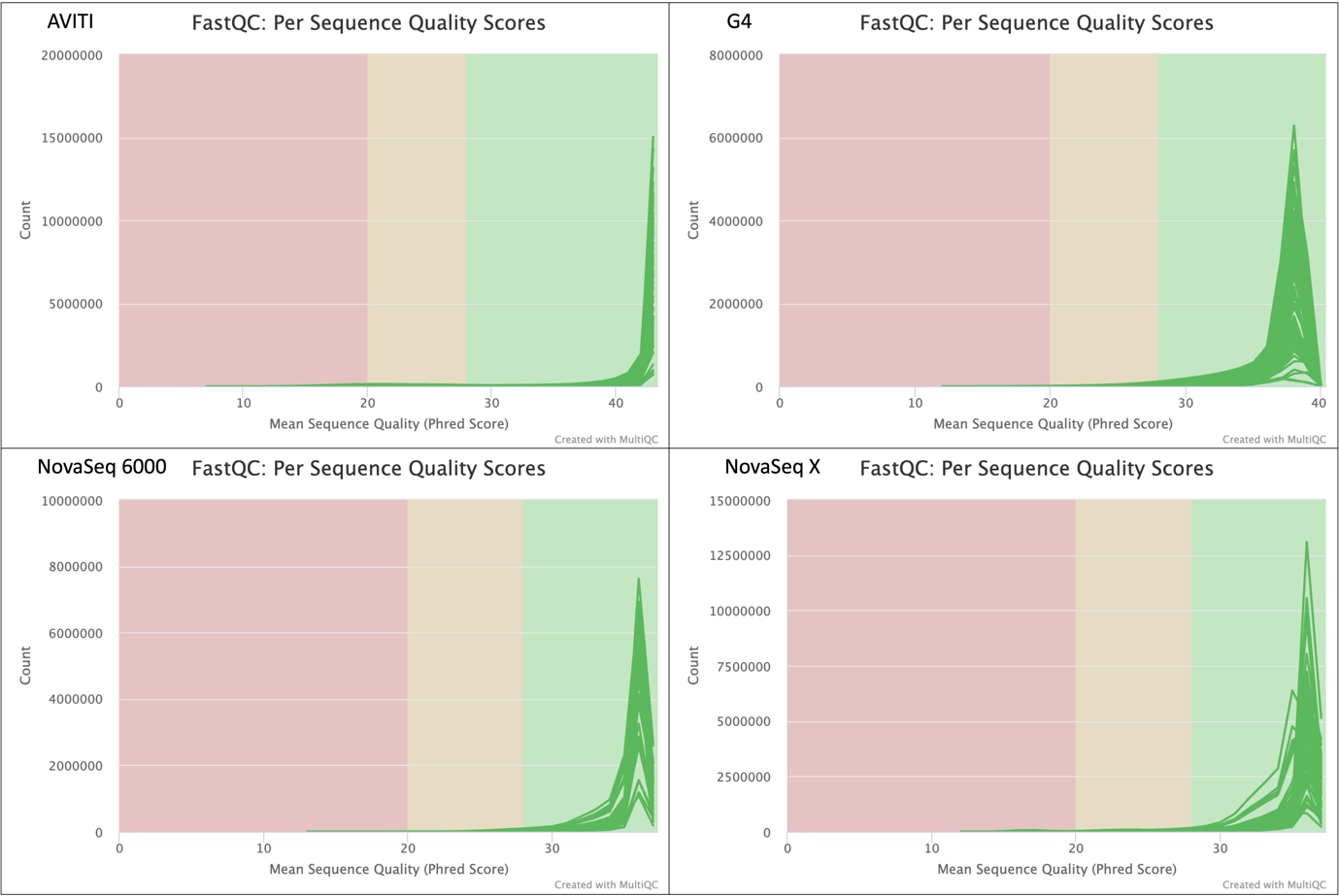

Supplementary Figure 1. Distribution of phred quality scores across all unsubsampled reads for all four sequencers, obtained from MultiQC reports.
